# Interferon signaling suppresses the unfolded protein response and induces cell death in hepatocytes accumulating hepatitis B surface antigen

**DOI:** 10.1101/2020.12.22.423938

**Authors:** Ian Baudi, Masanori Isogawa, Federica Moalli, Masaya Onishi, Keigo Kawashima, Yusuke Sato, Hideyoshi Harashima, Hiroyasu Ito, Tetsuya Ishikawa, Takaji Wakita, Matteo Iannacone, Yasuhito Tanaka

## Abstract

Virus infection, such as hepatitis B virus (HBV), often causes endoplasmic reticulum (ER) stress. The unfolded protein response (UPR) is counteractive machinery to ER stress, and the failure of UPR to cope with ER stress results in cell death. Mechanisms that regulate the balance between ER stress and UPR in HBV infection is poorly understood. Type 1 and type 2 interferons have been implicated in hepatic flares during chronic HBV infection. Here, we examined the interplay between ER stress, UPR, and IFNs using transgenic mice that express hepatitis B surface antigen (HBsAg) (HBs-Tg mice) and humanized-liver chimeric mice infected with HBV. IFNα causes severe and moderate liver injury in HBs-Tg mice and HBV infected chimeric mice, respectively. The degree of liver injury is directly correlated with HBsAg levels in the liver, and reduction of HBsAg in the transgenic mice alleviates IFNα mediated liver injury. Analyses of total gene expression and UPR biomarkers’ protein expression in the liver revealed that UPR is induced in HBs-Tg mice and HBV infected chimeric mice, indicating that HBsAg accumulation causes ER stress. Notably, IFNα administration transiently suppressed UPR biomarkers before liver injury without affecting intrahepatic HBsAg levels. Furthermore, UPR upregulation by glucose-regulated protein 78 (GRP78) suppression or low dose tunicamycin alleviated IFNα mediated liver injury. These results suggest that IFNα induces ER stress-associated cell death by reducing UPR. IFNγ uses the same mechanism to exert cytotoxicity to HBsAg accumulating hepatocytes. Collectively, our data reveal a previously unknown mechanism by which IFNs selectively induce cell death in virus-infected cells. This study also identifies UPR as a potential target for regulating ER stress-associated cell death.

**Author summary:** Hepatitis B virus (HBV) causes acute and chronic infections that kill over 600,000 people every year from severe hepatitis, liver cirrhosis, and cancer. Mechanisms of chronic liver injury remain largely unknown. Both type 1 and type 2 interferons (IFNs) have been implicated in hepatic flares during chronic HBV infection, although HBV per se is a poor IFN inducer. In addition, while IFNα, a type 1 IFN, used to be the first-line treatment for chronic hepatitis B (CHB) patients, adverse side effects, including hepatic flares, severely limit their therapeutic effectiveness. These clinical observations suggest a pathogenic role of IFNs in HBV infection. Here, we demonstrate that IFN-1s cause severe and moderate hepatitis in transgenic mice expressing hepatitis B surface antigen (HBs-Tg mice) and human hepatocyte chimeric mice infected with HBV, respectively. HBsAg accumulation appears to cause ER stress because a counteractive response to ER stress, namely, unfolded protein response (UPR), was induced in both HBs-Tg mice and HBV infected chimeric mice. Our results indicate that IFN-1s suppress UPR before causing liver injury. UPR was also suppressed by IFNγ. Induction of UPR in HBs-Tg mice before treatment with IFNα and IFNγ significantly alleviated liver injury. We suggest that IFNs exert cytotoxicity to ER stress accumulating cells by suppressing UPR.

## Introduction

Interferons (IFNs) play a critical role in host defense against pathogens, particularly viruses, by activating the expression of hundreds of genes that exert antiviral activity [1]. IFNs also cause immunopathology during viral infections [2,3]. The basis behind the IFN-mediated immunopathology has yet to be fully elucidated but appears to include phosphorylation of eukaryotic initiation factor-2α (eIF-2α), activation of RNase L, and induction of nitric oxide synthase (iNOS) [1,2]. Little is known about whether and how infected cells are preferentially sensitized to IFN-mediated cell death.

The endoplasmic reticulum (ER) is responsible for much of a cell’s protein synthesis and folding. An imbalance between the protein-folding load and the capacity of the ER causes unfolded or misfolded proteins to accumulate in the ER lumen, resulting in ER stress [4,5]. To cope with ER stress, a protective mechanism called the unfolded protein response (UPR) is activated to reduce protein synthesis and/or enhance degradation, folding, and secretion of the offending proteins [4,5]. UPR has been linked to various pathological states, including malignancies, neurodegenerative, storage, metabolic, and infectious diseases [6–11]. While unmitigated ER stress leads to cell death [12], it is unclear whether the concomitant UPR plays a pro- or anti-cell survival role in such cases. Maladaptation of UPR as the underlying pathogenic mechanism has been poorly studied, particularly in highly secretory organs such as the liver and pancreas that are prone to ER stress [13]. Although infections with several viruses have been reported to induce ER stress [14–19], the interplay between ER stress, UPR, and IFN signaling has not been adequately interrogated.

Hepatitis B virus (HBV) causes acute and chronic liver disease. Patients with chronic hepatitis B (CHB) often experience hepatic flares in association with hepatitis B surface antigen (HBsAg) accumulation [20,21]. Factors that cause hepatic flares in CHB patients are incompletely understood. IFNα, a type-1 IFN (IFN-1), is widely used as the first-line drug for the treatment of chronic hepatitis B (CHB) [22]. High serum IFNα levels have been reported in some CHB patients experiencing spontaneous hepatic flares [23] or antiviral therapy withdrawal-associated hepatic flares [24]. However, the molecular mechanisms behind the IFN-1 related liver injury largely remain obscure. In addition, CXCL10, an IFNγ inducible chemokine, has been associated with disease progression [25] as well as HBsAg clearance in CHB patients [26]. In this study, we examined the role of IFNs in liver injury associated with ER stress using transgenic mice that express HBsAg in the liver (HBs-Tg mice) and HBV-infected humanized-liver chimeric mice. Our data suggest that IFNs selectively causes cell death in hepatocytes under ER stress by perturbing UPR.

## Results

### Type-1 IFNs induce liver injury in association with HBsAg-retention

To investigate the role of IFN-1 signaling in liver injury associated with intrahepatic HBsAg accumulation, we used HBs-Tg mice (lineage 107-5D) that produce HBsAg [27]. Groups of 3-4 HBs-Tg mice or their non-transgenic littermates (WT) were intravenously injected with the IFN-1 inducer poly I:C or saline. Liver injury was monitored by measuring serum ALT activity (sALT) on days 1, 3, and 7 after treatment. As shown in Fig 1A, sALT was markedly elevated in the HBs-Tg mice, peaking on day 1 after poly I:C treatment, but not in the WT mice. We also examined whether poly I:C treatment could induce sALT elevation in HBV-replication-competent transgenic (HBV-Tg) mice (Lineage 1.3.32) [28]. As shown in Fig 1B, HBV-Tg mice readily secrete HBsAg, retaining approximately 100-fold less HBsAg in the liver compared with the HBs-Tg mice. Interestingly, no ALT elevation occurred in the HBV-Tg mice after poly I:C treatment (Fig 1C). Taken together, these results suggest that poly I:C-induced liver injury is associated with marked intrahepatic HBsAg accumulation.

**Fig 1.**
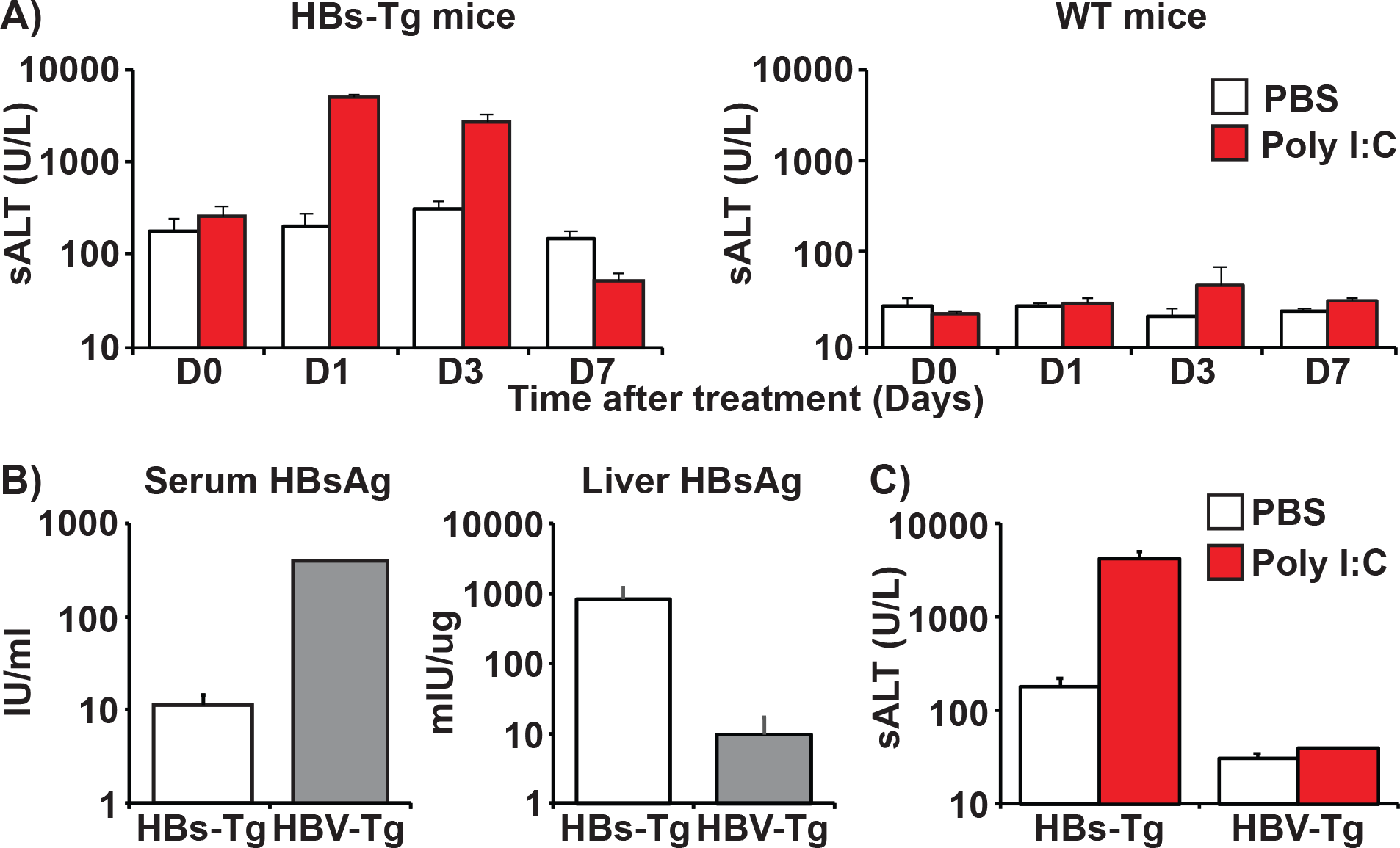
Poly I:C induces liver injury in association with HBsAg-retention. (A) Serum ALT levels measured on days 0, 1, 3, and 7 after PBS (white bars) or poly I:C (10 μg) (red bars) treatment in HBs-Tg mice (left graph) and WT mice (right graph). (B) Comparison of HBsAg expression levels between HBs-Tg (Lineage 107-5D: white bars) and HBV-Tg mice (Lineage 1.3.32: grey bars) in the serum (left graph) and the liver (right graph). (C) Serum ALT levels in HBs-Tg mice and HBV-Tg mice 24 hours after intravenous injection of PBS or poly I:C. Mean values +/− s.d of pooled data from 3 independent experiments are shown.

To examine the role of IFN-1 signaling in the poly I:C induced liver injury in HBs-Tg mice, groups of HBs-Tg mice (n=4) were treated with anti-IFNα/β receptor 1 (IFNαβR1) antibody that blocks IFN-1 signaling or a control antibody and then injected with poly I:C 24 hours later. The impact of IFN-1 signaling blockade was evaluated by monitoring sALT activity. As shown in Fig 2A, anti-IFNαβR1 antibody but not the control antibody treatment prevented sALT elevation, indicating that the liver injury induced after poly I:C treatment in HBs-Tg mice was dependent on IFN-1 signaling. To confirm that IFN-1 induces liver injury, groups of HBs-Tg mice (n=3) and control mice (n=3) were intravenously injected with 5 million units (MU)/kg of recombinant mouse IFNα, and sALT was monitored on days 1, 3, and 7. As expected, IFNα induced sALT elevation in the HBs-Tg mice, peaking on day 1 and receding towards baseline on day 3. No sALT elevation occurred in the normal mice (Fig 2B). Interestingly, liver immunohistochemical analysis showed almost no inflammatory cell infiltration at 24 hours despite severe sALT elevation, suggesting a sterile nature of IFNα mediated cell-death (Fig 2C). When apoptosis was examined by TUNEL staining at 16 hours after IFNα, a relatively small fraction of TUNEL positive hepatocytes were found scattered throughout the parenchyma (Fig 2D).

**Fig 2.**
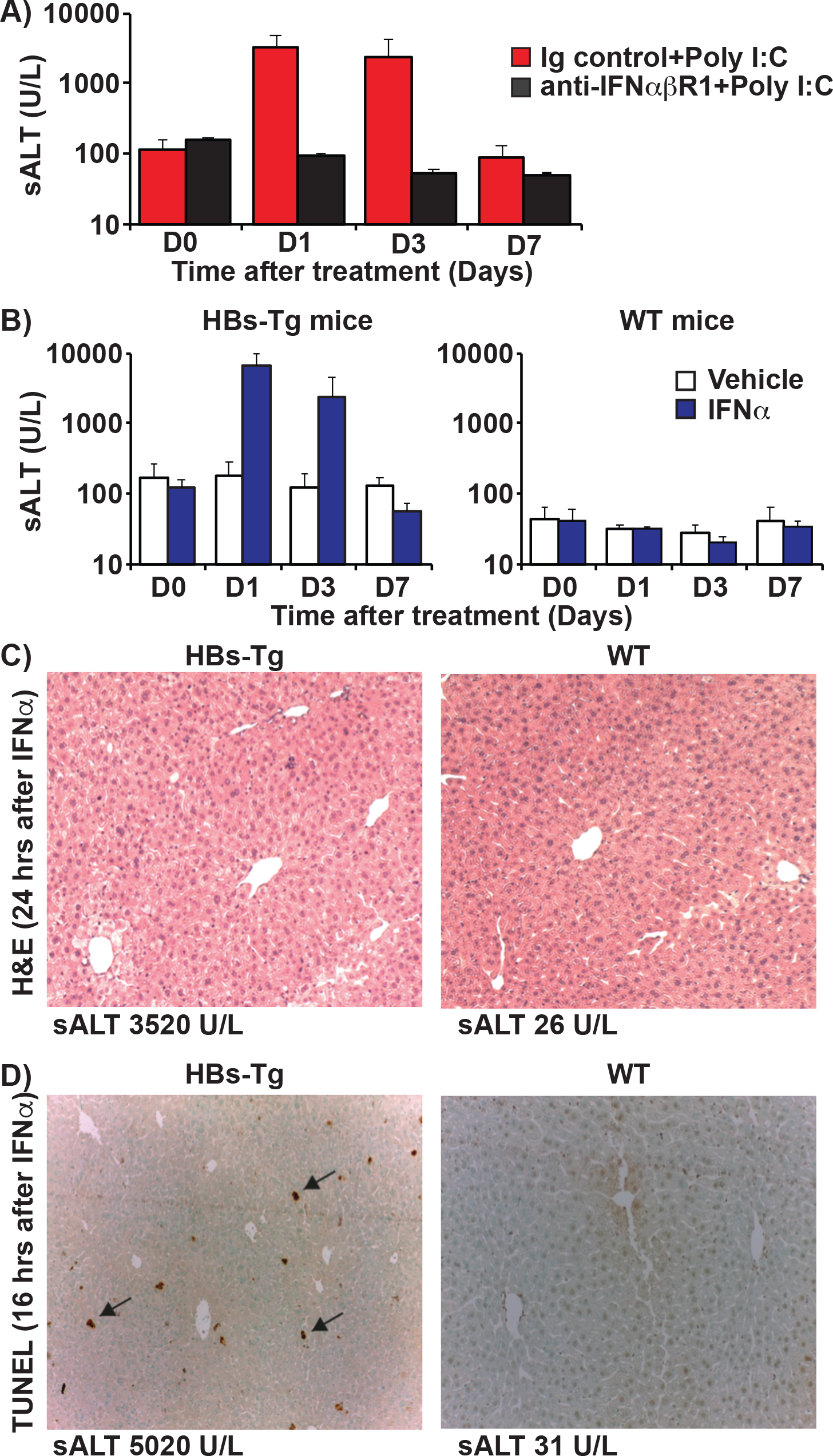
IFN-1s trigger HBsAg-associated liver injury in the absence of inflammatory cell infiltration. (A) Isotype control antibodies (250 μg: red bars) or anti-IFNαβR1 (250 μg: black bars) were intraperitoneally injected into HBs-Tg mice (n=3-4) 24 hours before poly I:C treatment. ALT levels on days 0, 1, 3, and 7 after poly I:C treatment are shown. (B) Serum ALT values in HBs-Tg (left graph) and WT mice (right graph) on days 0, 1, 3, and 7 after intravenous injection of PBS (white bars) or IFNα (5 million units (MU)/kg: blue bars). (C) Histological characteristics after IFNα injection in HBs-Tg mice. The panels show representative Hematoxylin and Eosin staining at 24 hours after IFNα. (D) TUNEL staining at 16 hours after IFNα treatment. The arrows show representative apoptotic cells in HBs-Tg mice.

To examine the impact of HBsAg reduction on IFN-mediated liver injury, HBs-Tg mice were injected by siRNA targeting HBV (siHBV) that were shown to reduce HBV mRNA in HBV-infected human hepatocyte chimeric mice [29,30] or control siRNA, and then intrahepatic HBsAg and HBV mRNA levels were determined 4 days later (S1A Fig). As shown in S1B Fig, siHBV had no discernable effect on HBsAg, although it strongly suppressed HBV mRNA expression (S1C Fig), indicating the stable nature of intracellular HBsAg. To induce hepatocyte turnover, we adoptively transferred HBsAg-specific T-cells that were shown to induce severe hepatitis in HBs-Tg mice [31] to groups of HBs-Tg mice (n=4) seven days before injecting siHBV or siControl. As shown in Fig 3B, HBsAg was significantly reduced on day 7 after siHBV treatment when hepatocyte turnover was previously induced. These mice were then injected intravenously with recombinant mouse IFNα on day 7 after siHBV treatment. A day after IFNα injection (day 8), serum ALT levels were significantly reduced in siHBV treated mice compared with their siControl treated littermates (sALT 425±103 vs. 4227±2533, p=0.027) (Fig 3C). These results indicate that IFNα mediated liver injury can be prevented by the reduction of intrahepatic HBsAg levels.

**Fig 3.**
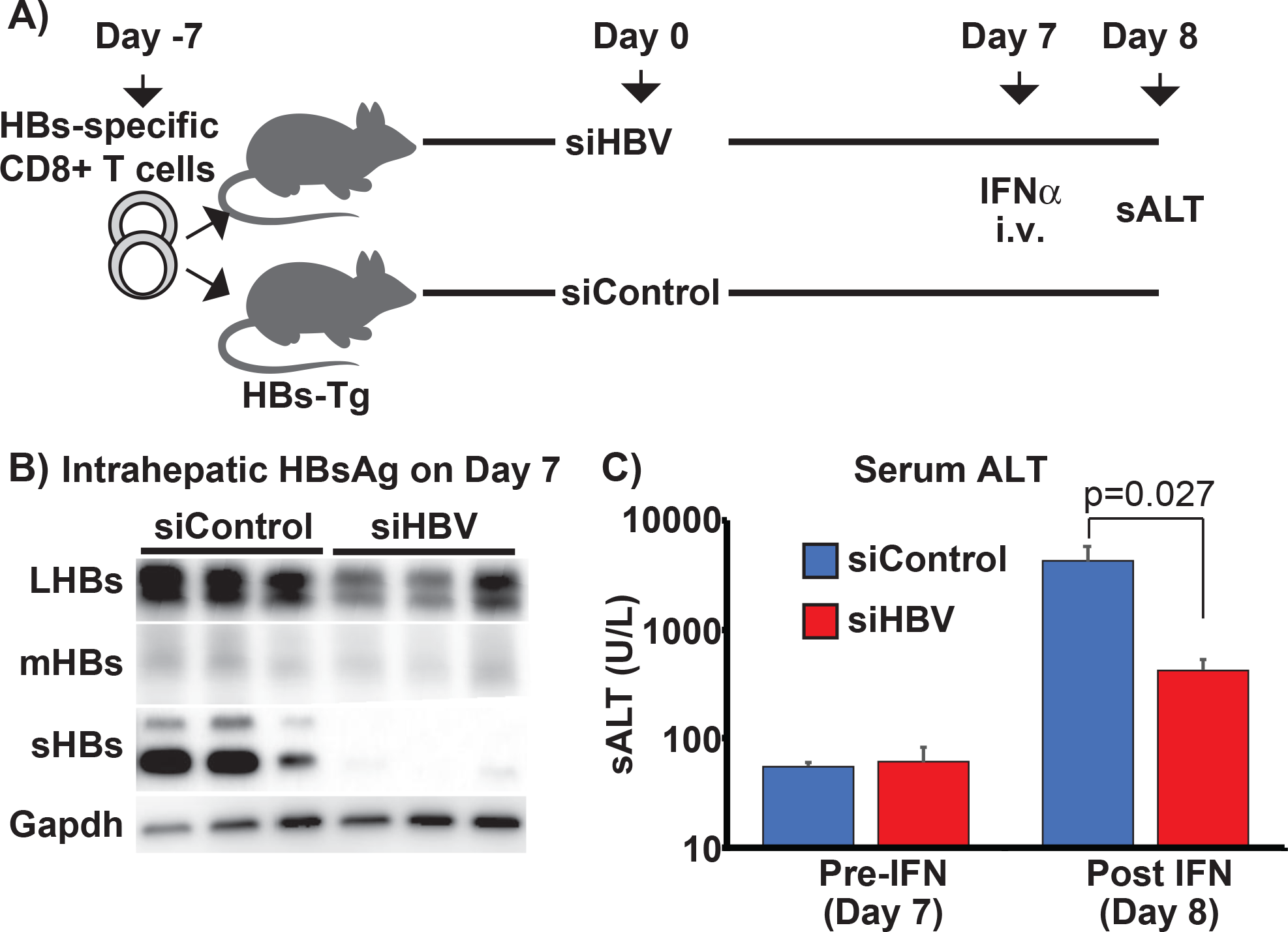
Reduction of intrahepatic HBsAg alleviates IFN mediated liver injury. (A) Experimental design. HBs-Tg mice were adoptively transferred with HBsAg-specific effector CD8 T cells on day 7 day before siRNA treatment. On day 7 after siRNA treatment, mice were intravenously injected with IFNα (5 MU/kg) and, on day 8, monitored for sALT levels. (B) Suppression of intrahepatic HBsAg expression before IFNα treatment (day 7 after siRNA treatment). (C) Serum ALT levels before (day 7) and after (day 8) IFNα treatment.

### IFN-1 signaling perturbs UPR in association with liver injury in HBs-Tg mice

To elucidate the molecular mechanism by which IFN-1 induces liver injury in HBs-Tg mice, we serially profiled the liver transcriptomes of both HBs-Tg and control mice treated with IFNα by microarray analyses. We especially searched for distinct genes whose expression correlated with the IFNα mediated liver injury in HBs-Tg mice. To do so, snap-frozen liver samples were obtained from groups of four HBs-Tg and control mice sacrificed prior to (0 hour) and 2, 4, 8, and 24 hours after treatment with 5 MU/kg of IFNα or vehicle. Liver injury progression was also monitored at these time points using sALT. As shown in Fig 4A, a significant sALT increase was observed in HBs-Tg mice from 8 hours post-IFN treatment and peaked at 24 hours (3758 ± 649 U/L). As expected, no sALT elevation occurred in the WT mice. Cluster and enrichment analyses were performed on the microarray data for the selected period prior and up to the onset of liver injury, i.e., 0, 2, 4, and 8-hour time-points. As shown in Figs 4B and 4C, six clusters showing distinct gene expression profiles were identified. Many ISGs could be found in TLR signaling and chemokine signaling pathways of Cluster 6 (Fig 4C), which were induced similarly between HBs-Tg and WT mice (Figs 4B and 4C). Cluster 3 genes were upregulated in HBs-Tg mice before IFN treatment, and as ISGs were induced, they were downregulated in both HBs-Tg and WT mice up to 4 hours (Fig 4B), suggesting a suppressive effect of ISGs on these genes. Interestingly, while the expression of these genes rebounded in WT mice, they remained downregulated in HBs-Tg mice at 8 hours after IFN treatment. Importantly, Cluster 3 includes genes related to protein processing in the endoplasmic reticulum (Fig 4C), raising the possibility that IFNα induced liver injury in HBs-Tg mice by modulating intrahepatic UPR. To test this, the correlation between liver injury, IFN-1 signaling, and UPR-related molecule expression was further examined. Intrahepatic induction of interferon-stimulated genes (ISGs), UPR biomarkers such as spliced X-box binding protein-1 (XBP1s) phosphorylated-PKR-like ER kinase (phos-PERK) and C/EBP homologous protein (CHOP) (Fig 4D) were assessed by western blot. As shown in S2A Fig, non-treated HBs-Tg mice showed higher baseline expression of XBP-1s, phos-PERK, and CHOP in association with moderate liver injury (sALT 100-300 U/L) than non-treated WT mice (S2B Fig), suggesting that HBsAg accumulation triggered UPR activation in association with mild spontaneous hepatitis. Interestingly, as shown in Fig 4E, the expression of XBP1-s and phos-PERK were transiently suppressed during ISG induction, in concordance with the mRNA levels represented in Cluster 3 of the microarray data (Figs 4B and 4C). As XBP1s and phos-PERK returned towards baseline levels, CHOP began to increase markedly in HBs-Tg mice from 8 hours and peaked at 24 hours in direct correlation with sALT. An ER chaperone molecule, glucose-regulated protein 78 (GRP78) was also upregulated in HBs-Tg mice at 24hours (Fig 4E), in association with peak liver injury (Fig 4A). The aforementioned changes were not observed in WT mice, although XBP1 expression was modestly reduced immediately after IFNα treatment (Fig 4E). These data suggest that IFN-signaling induces liver injury in HBs-Tg mice in association with UPR modulation.

**Fig 4.**
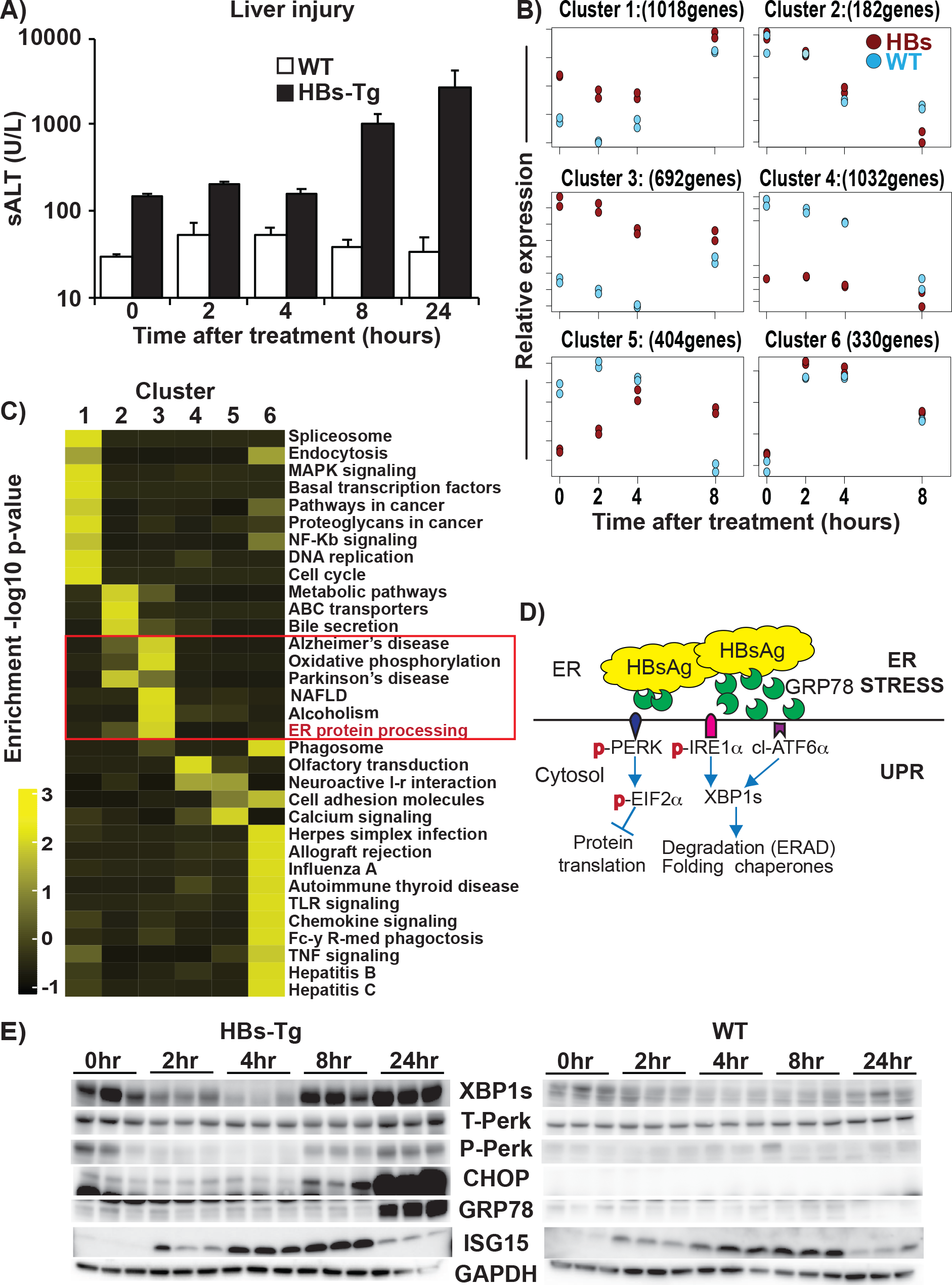
IFN-1 signaling perturbs UPR in association with liver injury in HBs-Tg mice. (A) Liver injury progression after IFNα treatment. The graph shows sALT levels in WT mice (white bars) and HBs-Tg mice (black bars) after IFNα treatment (5MU/kg). Mean values +/− s.d of pooled data from 3 independent experiments are shown. (B) Differentially expressed genes (DEGs) in the microarray data of HBs-Tg and WT mice after IFNα treatment. Each of the 6 graphs shows a distinct cluster identified using microarray time-course-specific maSigPro [32] software. (C) List of gene pathways selectively enriched in the 6 clusters. (D) A schematic diagram showing simplified unfolded response pathways. (E) Representative immunoblots showing UPR-biomarker protein expression in HBs-Tg mice (left panel) and WT mice (right panel) at specified time points after IFNα treatment.

### IFNα induces liver injury in HBV infected human chimeric mice in association with XBP1 suppression

To examine the effect of IFNα treatment on UPR in HBV infected human liver, we used human liver chimeric mice, i.e., uPA-SCID mice whose livers had been repopulated with human hepatocytes as previously described [33]. We first compared circulating and intracellular HBsAg levels between HBV-Tg mice, HBs-Tg mice, and chimeric mice infected with HBV for 6 weeks. As shown in S3 Fig, HBV infected chimeric mice secreted more HBsAg than HBV-Tg mice (>10-fold difference) and HBs-Tg mice (>100-fold difference) (S3A Fig) but still retained more HBsAg in the liver than 1.3.32 HBV-Tg mice (S3B Fig). Next, groups of nine chimeric mice were infected with HBV (genotype C isolate, C2_JPNAT, from a chronic HBV patient[33]), and after the establishment of persistent infection (6-8 weeks after HBV inoculation), the mice were intravenously administered with 25 ng/g of pegylated-human IFNα (PEG-hIFNα), and then sacrificed after 8 and 24 hours (n=3 per group) (Fig 5A) to examine the correlation between liver injury (sALT) (Fig 5B), IFN-1 signaling, and UPR-related molecules measured by quantitative-PCR (Fig 5C) and western blot (Fig 5D). Non-infected, non-treated control chimeric mice (n=2) were also sacrificed and included in the analyses. Before PEG-hIFNα treatment, HBV infected chimeric mice showed slightly higher sALT activity than untreated mice (Fig 5B, mean 268 vs. 130 U/L, p=0.001) in association with modest intrahepatic upregulation of UPR molecules including XBP1s and PERK (Fig 5C). Importantly, sALT levels increased in HBV infected chimeric mice at 8 (724 ± 159 U/L) and 24 hours (616 ± 26 U/L) after PEG-hIFNα treatment (Fig 5B) in association with an increased level of ISG15 expression and reduced levels of UPR-related molecules compared to non-treated HBV infected mice (Fig 5C and 5D). Intracellular HBsAg levels were mostly unchanged during the 24-hour period (Fig 5D), suggesting that the suppression of UPR-related molecules, including XBP1s, was not due to a reduction of HBV surface antigens. Collectively, these results indicate that bona fide HBV infection could induce UPR in human hepatocytes, and IFN-1 signaling induces liver injury in association with UPR suppression.

**Fig 5.**
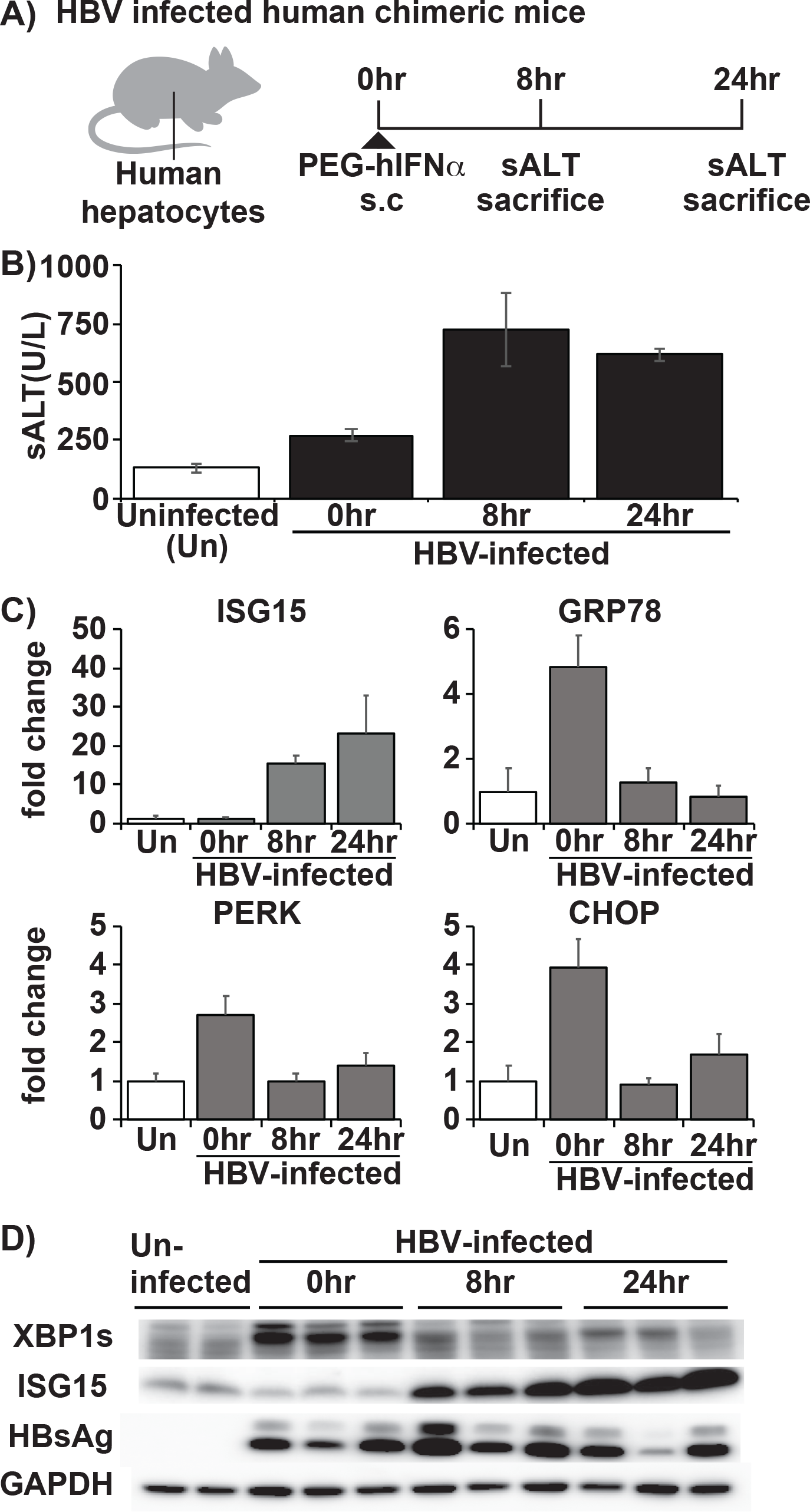
IFNα induces liver injury in HBV infected human chimeric mice in association with XBP1 suppression. (A) Experimental design. HBV infected mice were injected with 25 ng/g pegylated IFNα-2a and then sacrificed at specified time points (n=3 per time point). (B) Liver injury after IFN treatment in HBV infected chimeric mice. The graph shows serum ALT levels at baseline (0 h), 8, and 24 hours after IFN treatment (black bars) compared with non-infected non-treated chimeric mice (white bars). Data are shown as mean values +/− s.d. (C) Messenger RNA expression levels of UPR biomarkers after IFN treatment. Levels of UPR related molecule mRNA expression determined by quantitative PCR. (D) Association between XBP, ISG15, and HBV surface antigen (HBsAg) after IFN treatment. Representative immunoblots showing UPR biomarkers XBP1, ISG15, and HBsAg levels at 0, 8, and 24 hours after IFN treatment.

### IFNs exert direct cytotoxicity to HBsAg accumulating hepatocytes and downregulate UPR

To dissect the mechanism of the IFNα-mediated liver injury, we established a primary mouse hepatocyte (PMH) culture system amenable to knockdown experiments. Primary hepatocytes from both HBs-Tg and WT mice were treated with IFNα or medium, and then cytotoxicity and UPR-related protein expression were assessed 8, 24, and 72 hours later (Fig 6A). The degree of cytotoxicity was analyzed by calculating the percentage of specific cell death based on LDH activity in the supernatant as described in the Materials and Methods. As shown in Fig 6B, cytotoxicity was clearly observed at 72 hours after IFNα treatment in the HBs-Tg PMHs but not in the WT PMHs, indicating that IFNα is directly and specifically cytotoxic to HBsAg accumulating hepatocytes. Interestingly, UPR markers such as XBP1s, phosho-PERK, and CHOP expression were upregulated from 24 hours, and then diminished at 72 hours coinciding with ISG induction and cytotoxicity. Furthermore, GRP78 was upregulated at 72 hours in association with cytotoxicity. No UPR modulation was observed in the WT PMHs (Fig 6C). These in vitro data closely recapitulate our observations in vivo despite the slower temporal dynamics of ISG induction in vitro and lower baseline UPR in HBsAg PMHs. To determine whether the observed suppressive effect of IFNα on UPR was a general occurrence or restricted to the HBs-Tg model, we tested the impact of IFN-1 signaling on UPR that was chemically induced by thapsigargin in normal PMHs at 6 and 12 hours (Fig 6D). Interestingly, thapsigargin-induced Perk and CHOP but not XBP1s expression were clearly suppressed by IFNα treatment at both time points (Fig 6E), suggesting that UPR suppression by IFNα was in part a general phenomenon.

**Fig 6.**
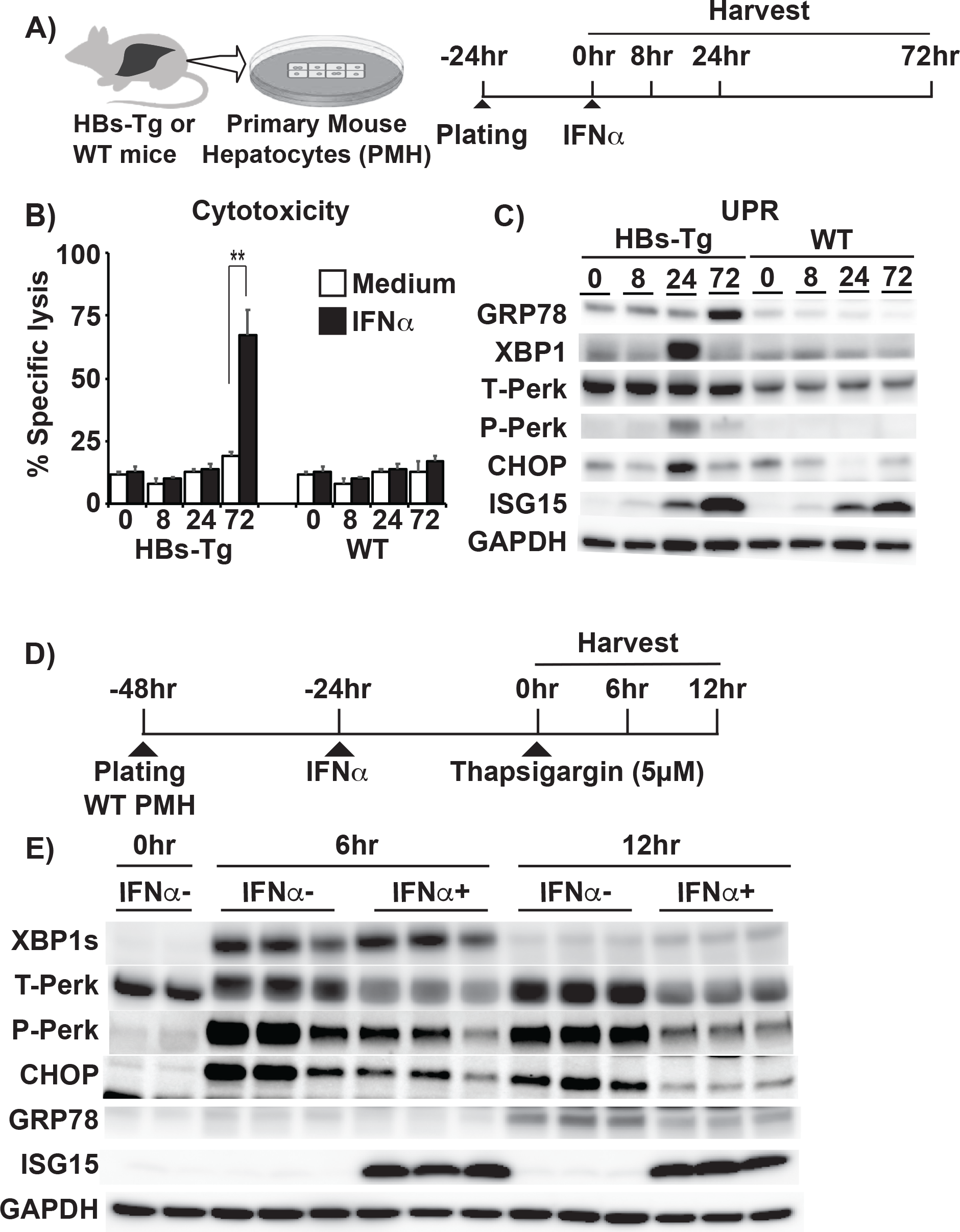
IFNα exerts direct cytotoxicity to HBsAg accumulating hepatocytes and downregulate UPR. (a-c) Effects of IFN signaling and UPR modulation on HBsAg accumulating hepatocytes in vitro. (A) Experimental design. (B) LDH levels in the supernatant of PMH culture were measured at indicated time points after adding medium (white bars) or IFNα (0.1MU/ml) (black bars). (C) The immunoblots of UPR related molecules and ISG15 at specified time points after IFNα treatment. (D, E) IFNα suppresses chemically induced UPR in vitro. (D) Experimental schema. (E) Representative immunoblots show the effect of IFNα on UPR related-protein levels at 0, 6, and 12 hours after treatment with thapsigargin.

We also tested the direct effect of IFNγ on hepatocytes accumulating HBsAg using the previously described in-vitro culture system (S4A Fig), because IFNγ has been reported to induce liver injury in mice that retain HBsAg in the liver [31,34]. As expected, IFNγ (10,000 U/ml) directly and specifically induced cell death in HBs-Tg PMHs (S4B Fig). Again, UPR was upregulated at 24 hours and then downregulated at 72 hours in association with cytotoxicity and GRP78 upregulation (S4C Fig), similar to IFNα.

### GRP78 suppression reduces IFN-induced hepatocytotoxicity by upregulating UPR

To determine the role of UPR related molecules in IFNα-mediated hepatocytotoxicity, CHOP, GRP78 and XBP1 were downregulated by transfecting HBs-Tg PMHs with target-specific siRNA or a scrambled siRNA control for 24 hours before addition of IFNα (Fig 7A). Cytotoxicity was assessed by measuring secreted LDH levels in culture supernatants 72 hours after IFNα addition. Surprisingly, suppression of GRP78 but not CHOP or XBP1 reduced LDH (Fig 7B) in association with upregulation of UPR related molecules, including PERK and XBP1s (Fig 7C). To determine whether the reduction in IFNα cytotoxicity by siGRP78 was due to the UPR upregulation, GRP78 and the key UPR molecule XBP1 were co-suppressed prior to IFNα treatment. Interestingly, the reduction of cytotoxicity by siGRP78 was lost when XBP1 was co-suppressed (Figs 7D and 7E), suggesting that UPR is required for the reduction of cytotoxicity by GRP78 suppression. These results suggest that robust UPR activation by GRP78 suppression rescues HBsAg accumulating hepatocytes from the cytotoxic effect of IFNα.

**Fig 7.**
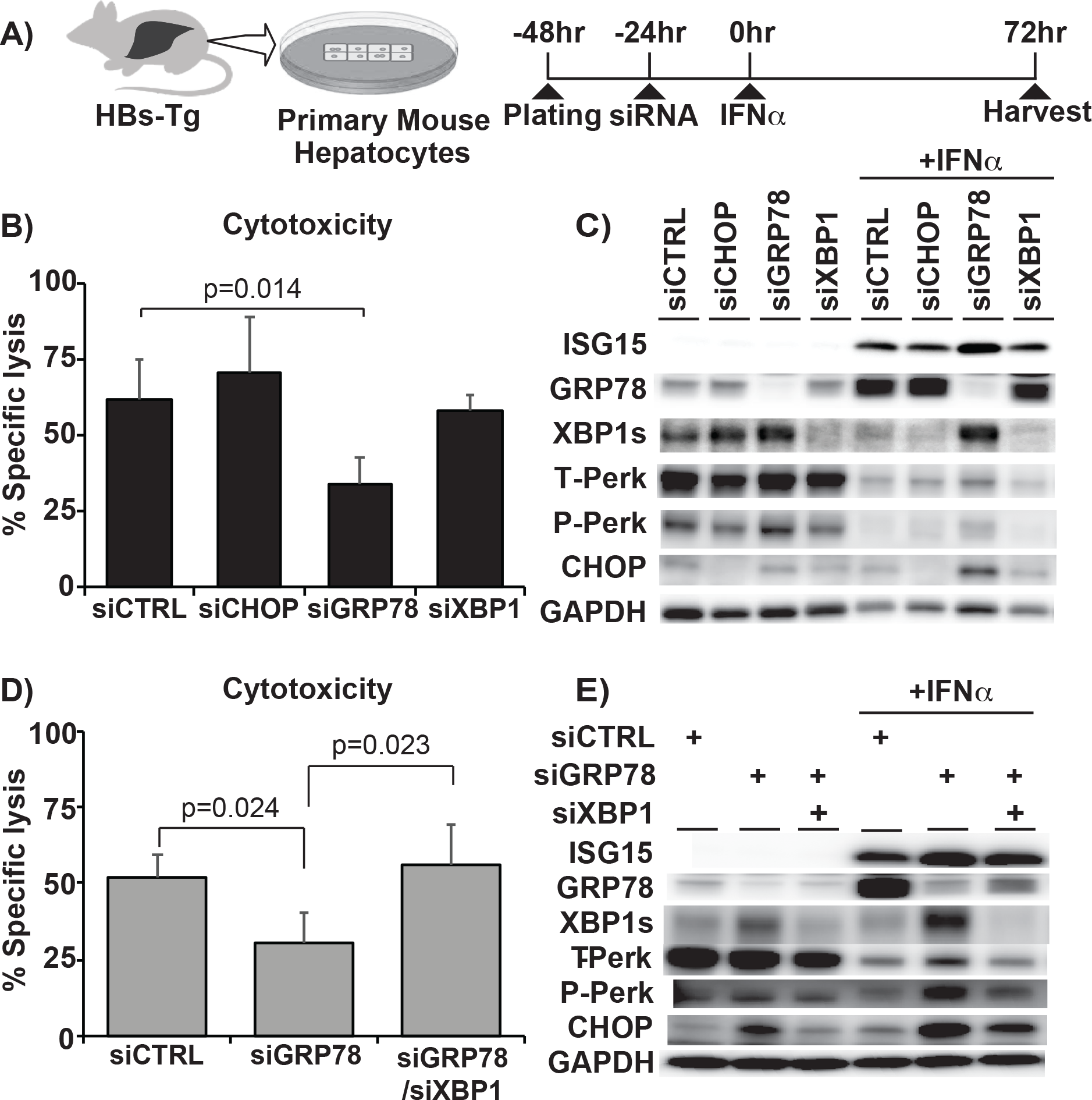
GRP78 suppression reduces IFNα-induced hepatocytotoxicity in association with UPR upregulation. (A) Experimental design. Small interfering RNA (siRNA) targeting CHOP, GRP78, XBP1, or control scramble siRNA (siControl) (15 µM) were transfected to primary mouse hepatocytes (PMHs) from HBs-Tg and WT mice before IFNα treatment. (B) LDH levels in the supernatant of cultured HBs-Tg PMHs after IFNα treatment following knockdown of each UPR-related molecule by specific siRNA. (C) Representative immunoblots showing UPR-related protein levels in the absence/presence of IFNα after indicated specific target downregulation by siRNA. (D) LDH levels in the supernatant of cultured HBs-Tg PMHs after IFN treatment following control, GRP78, and GRP78/XBP1 co-suppression by siRNA. (E) UPR-related protein levels in the absence/presence of IFNα in siControl, siGRP78, and siGRP78/siXBP1 suppressed HBs-Tg PMHs. Mean values +/− s.d of pooled data from 2 independent experiments are shown.

### UPR upregulation alleviates IFN-induced liver injury in HBs-Tg mice

To determine whether GRP78 suppression could alleviate liver injury in HBs-Tg mice, we intravenously injected siRNA targeting GRP78 (siGRP78), or control siRNA, into groups of 3-4 HBs-Tg mice 4 days before IFNα treatment (Fig 8A). As shown in Fig 8B, siGRP78 treated mice exhibited 3-5-fold lower sALT levels after IFNα treatment compared to control siRNA treated mice (mean 7110 ± 42 U/L vs. 1610 ± 794 U/L, p=0.003). These data suggest that robust UPR induction prevents IFN-induced liver injury in HBs-Tg mice. Therefore, the impact of UPR augmentation on IFN-mediated liver injury was tested using a chemical UPR inducer, tunicamycin. Groups of HBs-Tg and WT mice (n=3) were intraperitoneally injected with a low dose of tunicamycin (0.1 mg/kg) or saline. Significant UPR upregulation could be seen 4 hours after tunicamycin treatment (Fig 8C), at which time point IFNα (5 MU/kg) or 50 ng α-Galactocsylceramide (αGal; an IFNγ inducer) was injected into these mice. The levels of sALT activity and intrahepatic UPR-related molecule expression were measured 24 hours after IFNα or αGal treatment. As shown in Fig 8D, sALT elevation was almost completely blocked in the tunicamycin pre-treated animals compared to the vehicle-treated control group after IFNα (3665±500 U/L vs. 207±73 U/L, p<0.001). Tunicamycin treatment also suppressed IFNγ-mediated liver injury in HBs-Tg mice as tunicamycin treated mice showed up to a 6-fold reduction in sALT levels compared to the controls after αGal injection (Fig 8E; 12020±1100 U/L vs. 1940±388 U/L, p<0.001). These data indicate that UPR augmentation rescues HBsAg accumulating cells from the cytolytic effect of type I and type II IFNs. The data also suggest that IFNα and IFNγ utilize a similar molecular mechanism to induce cell death in HBsAg accumulating hepatocytes (Fig 8F).

**Fig 8.**
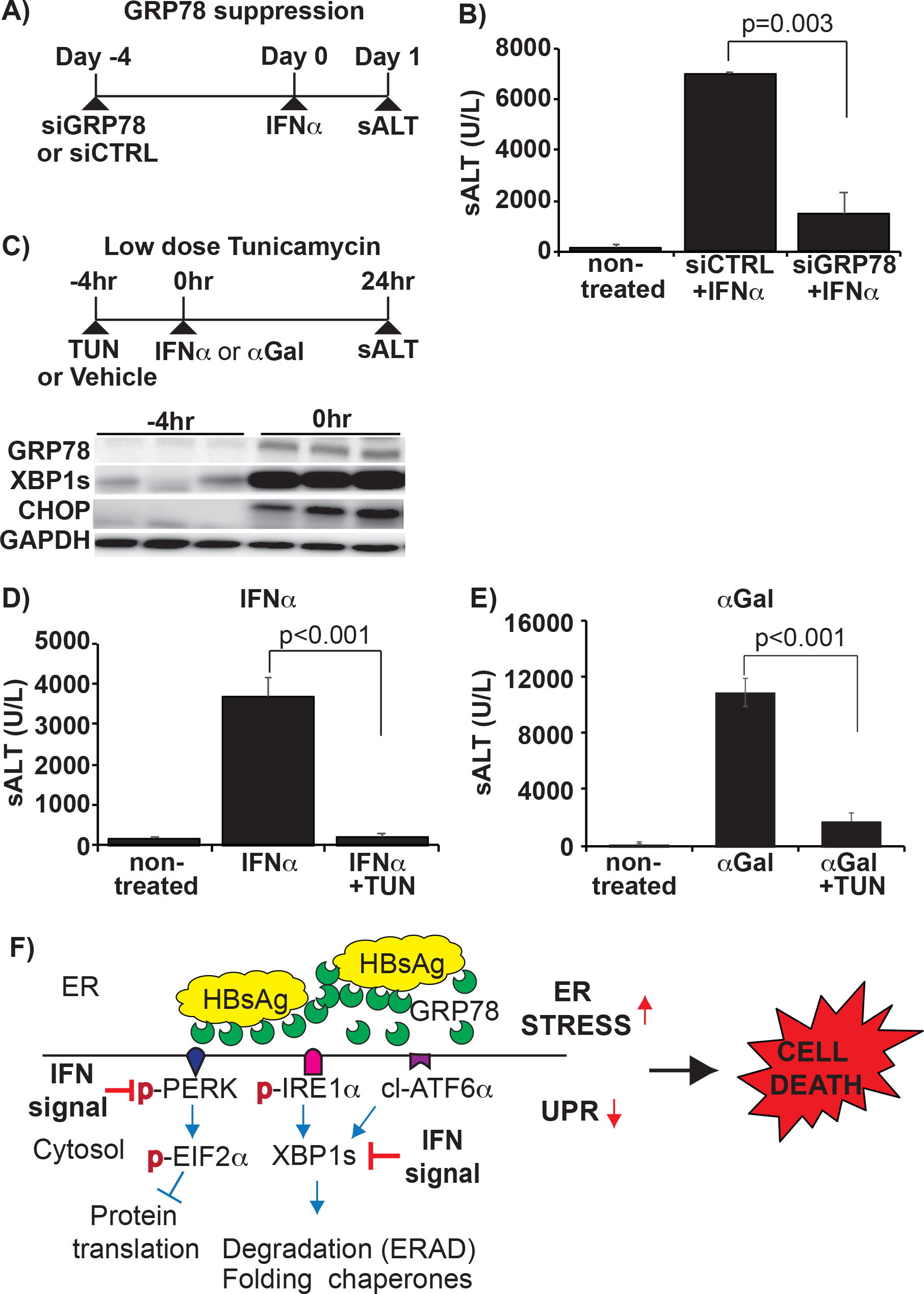
UPR upregulation alleviates IFN-induced liver injury in HBs-Tg mice. (A) Experimental design to test the effect of GRP78 suppression on IFN-induced liver injury. (B) sALT levels at 24 hours after IFNα injection in siGRP78 treated HBs-Tg mice compared with controls. (C) Experimental design to test the effect of tunicamycin administration (0.1mg/kg) on IFN-induced liver injury. IFNα (5 MU/kg) or α-Galactocsylceramide (αGal) (50ng/mouse) were intravenously injected to HBs-Tg mice at 4 hours after tunicamycin administration. The immunoblots below show the intrahepatic level of UPR markers at indicated time points. (D) sALT levels at 24 hours after IFNα injection in tunicamycin (TUN) treated HBs-Tg mice compared with controls. (E) sALT levels at 24 hours after αGal injection in tunicamycin and vehicle-treated HBs-Tg mice. Mean values +/− s.d of pooled data from 3 independent experiments are shown. (F) A schema depicting our hypotheses on the effect of IFN-1 signaling on UPR to induce ER stress-related cell death in HBsAg accumulating hepatocytes.

## Discussion

In this study, we examined whether and how IFNs induce liver injury in association with HBsAg accumulation in the liver. IFNα directly and specifically induced cell death in HBsAg accumulating hepatocytes in association with suppression of the pro-survival UPR. IFNγ appears to use the same mechanism to cause cellular damage. Importantly, UPR augmentation significantly reduced the cytolytic effect of IFNs on HBsAg accumulating hepatocytes. Our data highlights an as yet unknown characteristic of the IFN signaling-UPR axis that potentially presents important targets for regulating ER-stress associated cell death.

The HBV envelope consists of three closely related envelope proteins: small (S), middle (M), and large (L), all of which have identical C-terminal ends [35]. The large envelope protein (LHBs) is filamentous and tends to accumulate in the ER [36]. Lineage 107-5D transgenic mice used in this study produce LHBs predominantly, and were shown to be exquisitely sensitive to IFNγ [31,34,37,38]. IFNγ was also shown to induce cell death in oligodendrocytes that accumulate MHC-1 heavy chains [39]. However, little is known about the mechanism by which IFNγ induces cell death in ER stress-accumulating cells. In contrast, the impact of type I IFNs on ER stress-accumulating cells had not been examined. Due to intracellular pattern recognition receptors (PRRs) such as RIG-I, IFNα can be specifically induced by virus-infected cells [3]. Less understood was whether and how IFNα specifically targets virus-infected cells to exert its antiviral activity. Although many viruses are shown to induce ER stress and UPR [14–19], the interaction between IFNs, ER stress, and UPR remains largely unexplored. To our knowledge, this is the first study that demonstrates the direct and specific cytotoxicity of IFNα on ER stress-accumulating cells.

Importantly, the suppressive effect of IFN on UPR was evident not only in HBs-Tg mice (Fig 4) and HBV infected chimeric mice (Fig 5), but also on chemically induced UPR (Fig 6D and 6E). IFNα suppressed the expression of phosphorylated and unphosphorylated PERK, which is a key UPR molecule presumed to suppress protein synthesis [40]. In HBs-Tg mice and HBV infected chimeric mice, IFNα also suppressed the expression of spliced XBP1. XBP1 is a key transcription factor that binds to unfolded protein response elements (UPRE) found in several genes encoding molecular chaperones and ER-associated degradation (ERAD) [41]. IFNα did not suppress XBP1 in thapsigargin-treated PMHs. Of note, thapsigargin treatment did not increase LDH in the supernatant, indicating that the nature of ER stress caused by HBs-Tg mice and thapsigargin is very different. Previous studies have shown that hepatic deficiency of PERK [42] or XBP1 [43,44] renders hepatocytes hypersusceptible to ER stress-related cell death and disease. It is currently unclear whether the suppression of these molecules is the primary cause triggering the cascade of cell death after IFN treatment. Knockdown of the key UPR related molecules such as XBP1 and PERK individually by siRNA did not induce cell death in the absence of IFNs. It is possible that the simultaneous suppression of several UPR molecules or other unknown factors induced by IFNs is required to initiate the cytolytic program. Alternatively, the suppression of UPR by IFN signaling may be cytolytic only when GRP78 is highly upregulated. Regardless, the current study demonstrated the detrimental effect of UPR suppression on HBsAg accumulating hepatocytes.

GRP78 was upregulated after IFN treatment selectively in HBs-Tg mice, and its upregulation was clearly associated with liver injury. Because GRP78 regulates the UPR through direct interaction with ER stress sensors [45], it is reasonable that the upregulation of GRP78 results in reduced UPR. Paradoxically, the main function of GRP78 is thought to facilitate protein folding to reduce ER stress, and its expression is also induced by UPR stimulation [46]. Ample evidence suggests the critical role of GRP78 in cancer cell survival and proliferation. Site-specific deletion of GRP78 in the prostate epithelium suppresses prostate tumorigenesis [47], and knockdown of GRP78 sensitizes various cancer cells to chemotoxic, anti-hormonal, DNA damaging, and anti-angiogenesis agents [48]. In stark contrast to these reports, more recent studies point to a pro-apoptotic role of GRP78 translocated to the cell surface. Ligation of a proapoptotic protein, prostate apoptosis response-4 (Par-4), to GRP78 on cell surface induces apoptosis [49]. Experiments are currently underway to test whether IFNs induce GRP78 translocation to the surface of HBsAg accumulating hepatocytes.

CHOP does not seem to play a significant role in IFNα mediated cell death associated with HBsAg accumulation in our setting (Fig 7B), although it was highly induced in direct correlation with ALT elevation (Fig 4E). This observation appears to contradict the widely accepted notion that CHOP sensitizes cells to ER stress-mediated death. For example, cells lacking CHOP are significantly protected from the lethal consequences of ER stress [50,51]. However, mouse embryonic fibroblasts (MEFs) derived from CHOP-knockout mice exhibit only partial resistance to ER stress-driven apoptosis [50]. Furthermore, liver-specific CHOP knockdown had no impact on liver damage associated with urokinase plasminogen activator (uPA) accumulation in the ER [52]. Thus, it is possible that the role of CHOP depends on the organs and nature of ER stress.

Clinically, we do not know to what extent the observed phenomenon contributes to the liver disease in HBV infected patients as HBV is a poor IFN-1-inducer [53]. However, liver damage is one of the most common side effects of IFNα treatment [54]. In addition, accumulating evidence suggests that ALT flares during chronic HBV infection are associated with increases in serum IFN-1s [23,24]. Liver injury is often severe in CHB patients superinfected with hepatitis C or hepatitis D viruses [55,56], both of which induce IFN-1s. While activated NK, NKT, and T cells have been assumed responsible for the liver injury associated with IFNs [23,24], the current study newly identifies UPR downregulation as a potential mechanism of IFN-mediated hepatoxicity. One may argue that the levels of HBsAg accumulation in the HBs-Tg mice cannot be attained in HBV infected patients. However, some chronic hepatitis B patients, particularly those carrying HBV PreS1/PreS2 mutants [57–59], exhibit ground glass hepatocytes (GGHs), a manifestation of HBsAg accumulation in the ER, and tend to develop hepatic flares. Immune suppressed CHB patients occasionally experience fibrosing cholestatic hepatitis, a severe form of hepatic flares, and exhibit GGHs [20,21]. GGHs have been reported even in HBV-infected chimeric mice in association with very high levels of intrahepatic HBV proteins and also mild hepatitis [60]. It is important to point out that UPR was induced in HBV infected human hepatocytes in chimeric mice, and UPR downregulation was observed in these mice after IFNα treatment, indicating that the similar events observed in HBs-Tg mice could occur during bona fide HBV infection.

In conclusion, the results described herein suggest the previously unappreciated mechanism by which IFNs selectively induces cell death in ER stress accumulating cells. This mechanism may have evolved to selectively eliminate stressed cells due to virus infections or other causes. On the other hand, the same mechanism potentially induces chronic inflammation. Further studies are warranted to determine whether the IFN-UPR axis contributes to the development of other ER stress-associated chronic inflammatory diseases, such as alcoholic and non-alcoholic steatohepatitis, and neurodegenerative disorders.

## Materials and Methods

### Ethics statement

All experiments involving mice were performed in the Center for Experimental Animal Science at Nagoya City University, following a protocol approved by the Institutional Animal Care and Use Committee of the Nagoya City University Graduate School of Medical Sciences (approved number: H30M_45).

### Mouse models and treatments

Mouse care and experiments were performed at the Nagoya City University Center for Experimental Animal Science following a protocol approved by the Institutional Animal Care and Use Committee. Ten to twelve-week-old male mice were used in all the experiments. HBV transgenic mice (Lineage 1.3.32) and HBs-Tg mice of the lineage 107-5D used in this study were kindly provided by Dr. Francis V. Chisari. HBs-Tg mice produce filamentous HBs proteins under the control of the albumin promoter, as previously described[27]. HBV-Tg mice produce infectious Dane particles and HBV subviral particles as described previously [28]. Human hepatocyte chimeric mice were generated by repopulating the livers of severe combined immunodeficient mice transgenic for the urokinase-type plasminogen activator gene (uPA^+/+^/SCID^+/+^ mice) with human hepatocytes, and purchased from Phoenix Bio Co., Ltd, (Hiroshima Japan). Chimeric mice were infected with HBV as previously described ^15,^ and 3-4 mice/group subcutaneously received 25ng/g of pegylated-human interferon α (Peg-hIFNα-2a) (Hoffmann La Roche, Basel, Switzerland). To activate IFN-1 signaling, poly I:C (Sigma) (10 μg/mouse) was injected intravenously. To block type-1 IFN signaling, 250µg anti-IFNαβR1 antibody (clone MAR1-5A3) or isotype control (IgG1) (BioXcell, Lebanon, NH, USA) were injected intraperitoneally. To test the effect of IFNα, 5 million units per kg recombinant mouse IFNα (Miltenyi Biotec) were injected intravenously. To upregulate UPR, 0.1mg/kg tunicamycin was injected intraperitoneally (Merck).

### Serum ALT and HBsAg analyses

Serum ALT was measured using the Dri-Chem 3500 analyzer according to the manufacturer’s instructions (Fuji, Tokyo, Japan). Serum HBsAg and intrahepatic HBsAg were measured by a chemiluminescent enzyme immunoassay (CLEIA) using a LumipulseG1200 analyzer (Fujirebio, Tokyo, Japan), as previously described [33].

### Detection of DNA fragmentation by TUNEL method

Mouse livers were perfused with PBS, harvested in Zn-formalin and transferred into 70% ethanol 24 hours later. Tissue was then processed, embedded in paraffin and stained as previously described [61]. In situ apoptosis detection was carried out by using a Terminal deoxynucleotidyl transferase dUTP nick end labelling (TUNEL) apoptosis kit, according to the manufacturer’s instructions (ab206386; Abcam Biotechnology, Cambridge, UK). The samples were counterstained with Mayer’s hematoxylin for the morphological evaluation and characterization of normal and apoptotic cells.

### Adoptive HBsAg-specific T cell transfer and In vivo HBsAg suppression

HBsAg-specific CD8+ T cells, derived from TCR transgenic mice (Env-28 specific, Lineage: 6C2.16) [62], were simulated in-vitro for one-week as previously described [31]. These cells were intravenously injected into groups of HBs-Tg mice at a dose of 5 ×10^6^ cells/mouse. Groups of HBs-Tg mice were intravenously injected with a single dose (5mg/kg) cocktail of HBV specific siRNA (si75, si251, si1803)[29,30,63] or Control siRNA complexed with a pH-sensitive multifunctional envelope-type nanodevice (MEND) [29].

### Immunoblots

Whole-cell extracts were obtained from liver tissue or cell pellets lysed in buffer (0.1%, sodium dodecyl sulfate, 0.1% sodium deoxycholate, 1% IGEPAL) supplemented with Protease and Phosphatase Inhibitor cocktails (Roche). Protein extracts were separated by SDS-polyacrylamide gel electrophoresis then transferred onto polyvinylidene difluoride (PVDF) membranes (Millipore, Temecula, CA). Primary antibodies and secondary antibodies were used according to manufacturers’ instructions. Primary antibodies used are GRP78 (#3177), CHOP (#2895), XBP1s (#12782), phosphor-PERK (#3179) PERK (#3192), ISG15 (#2743) IRE1α (#3294), Stat-1 (#9172), phospho-Stat-1 (#9167) (all from Cell Signaling Technologies), GAPDH (abcam#8245), HBsAg (#5124A, Tokumen, Japan) phospho-IRE1α (Novus #NB100-2323) Detection was performed using the Immobilon Western Chemiluminescent HRP Substrate (Millipore) and image capture was done using the A1680 imaging system (GE Healthcare).

### RNA extraction and gene expression analyses

RNA was isolated from snap-frozen liver tissue obtained at selected time points using Isogen (Nippon Gene, Tokyo, Japan) according to the manufacturer’s instructions. For Microarray analyses, 2 biological replicates for each selected time point were analyzed. Briefly, from 100ng total RNA, complementary RNA (cRNA) was prepared using the Low Input Quick-Amp Labeling Kit, one color Cy3 protocol (Agilent Technologies, Santa Clara, CA, USA). Purified labeled-cRNA and controls (Agilent One Color Spike-In Kit) were hybridized to Agilent SurePrint G3 Mouse Gene Expression v2.0 Microarray Chips. Detection, data extraction, and pre-analysis were performed using a G2505C Agilent microarray scanner, Feature Extraction v10.10.1.1, and GeneSpring GX v14.8.0 software. Genes showing differential kinetics between IFN treated or non-treated HBsAg and WT samples over the period leading up to the onset of liver injury in HBs-Tg mice, i.e., 0, 2, 4, and 8 hour time points, were identified using the time-course specific R program maSigPro [32]. Briefly, maSigPro assesses the significance of the global model (i.e., if there are significant differences with respect to time or treatment) and of each variable (i.e., which specific time or treatment change is present) by fitting a regression model, considering time as a continuous variable and creating specific variables for each treatment, thereby adjusting a temporal profile for each time course. Genes significantly changed were selected and divided into clusters of similar profiles for visualization of results. Microarray data have been deposited into the NCBI Gene Expression Omnibus Repository (Accession # GSE 138916)

For microarray data validation and specific target gene expression analysis, quantitative real-time polymerase chain reaction (qRT-PCR) was performed using the StepOnePlus Real-Time PCR System (Applied Biosystems, Foster City, CA). From 2μg total RNA, complementary DNA (cDNA) was synthesized using the High Capacity RNA-to-cDNA Kit (Applied Biosystems). TaqMan Gene Expression Assay primer-probe sets used include; GRP78 (Mm00517690_g1, Hs00607129_gH), CHOP (Mm01135937_g1, Hs03834620_s1), XBP1s (Custom), PERK (Mm00438700_m1, Hs00984006_m1), GAPDH (Mm99999915_g1, Hs02758991_g1), IRE1α (Mm00470233_m1), ISG15 (Mm01705338_s1, Hs01921425_s1), IRF1(Hs00971960_m1) (all from Applied Biosystems). All qPCR data were normalized to GAPDH.

### Isolation, culture, and treatment of primary mouse hepatocytes

Primary mouse hepatocytes were isolated from both HBs-Tg and their wildtype littermates (controls) using a two-step protocol as previously described [62] with some modification. Briefly, the liver was perfused with Liver Perfusion Medium (Gibco #17701) for 4 minutes at a flow rate of 5ml/min followed by 0.8mg/ml Type 1 Collagenase (Worthington, UK) in Dulbecco Minimum Essential Medium (Gibco # 11965-092) for 8-12 minutes at 5ml/min. PMHs were cultured on Collagen 1 Biocoat 6-well plates (BD Biosciences) at a seeding density of 1×10^5^cells/cm^2^ in Hepatocyte Growth Medium with 2% DMSO. Thapsigargin (5µM, Fujifilm-Wako, Osaka, Japan) was added to IFNα-treated WT-PMHs that were then harvested after 6 and 12 hours. Recombinant mouse IFNα was added at 0.1MU/ml. Recombinant mouse IFNγ was added at 0.01MU/ml per well.

### LDH activity assay

Cytotoxicity was measured by the release of lactate dehydrogenase (LDH) into the culture media 72 hours after the addition of IFNα (final concentration, 0.1MU/ml). Collected supernatants were assayed using the Dri-Chem analyzer according to the manufacturer’s instructions (Fuji, Tokyo, Japan). Data are presented as percentage specific cell lysis, calculated as the ratio of the experimental LDH release (minus spontaneous release LDH) to the maximum LDH release using 1% Triton-X (minus spontaneous release LDH) for each plate.

% specific lysis = LDH [(experimental-spontaneous)/maximum lysis – spontaneous] ×100

### Knockdown of UPR-related target genes

To suppress UPR related molecules in vivo and in vitro, the Invivofectamine 3.0 and Lipofectamine RNAiMAx transfection reagents were used respectively, together with siRNA for the following targets; Control (#1), GRP78 (s607083, s67085), XBP1 (s76114), CHOP (s64888), all according to manufacturer’s instructions (all from Thermofisher Scientific).

### Statistics

Student’s t-test and One-Way ANOVA were performed accordingly. Microarray data bioinformatic analyses were performed as previously described [32]. Data are depicted as the mean ± SD, and P values < 0.05 were considered significant.

## Acknowledgements

The authors are grateful to Dr. Francis V. Chisari (Scripps Research Institute) who produced and provided the HBV and HBs-Tg mice used in this study and for his critical reading of the manuscript. We also thank Takayo Takagi, Mayumi Hojo, and Kyoko Ito (Nagoya City University, Japan) for their technical assistance.

## Author contributions

Conceptualization: Masanori Isogawa

Data Curation: Ian Baudi, Masanori Isogawa, Federica Moalli, Masaya Onishi, and Keigo Kawashima

Formal analysis: Masaya Onishi

Funding Acquisition: Masanori Isogawa, Takaji Wakita, Matteo Iannacone, and Yasuhito Tanaka

Investigation: Ian Baudi, Masanori Isogawa, Federica Moalli, and Keigo Kawashima

Methodology: Ian Baudi, Masanori Isogawa, Federica Moalli, and Matteo Iannacone.

Resources: Yusuke Sato, Hideyoshi Harashima, Hiroyasu Ito, Tetsuya Ishikawa

Supervision: Masanori Isogawa, Yasuhito Tanaka

Writing – original draft: Ian Baudi, and Masanori Isogawa

Writing – review & editing: Ian Baudi, Masanori Isogawa, Federica Moalli, Masaya Onishi, Keigo Kawashima, Yusuke Sato, Hideyoshi Harashima, Hiroyasu Ito, Tetsuya Ishikawa, Takaji Wakita, Matteo Iannacone, and Yasuhito Tanaka

## Supporting information

**S1 Fig.**
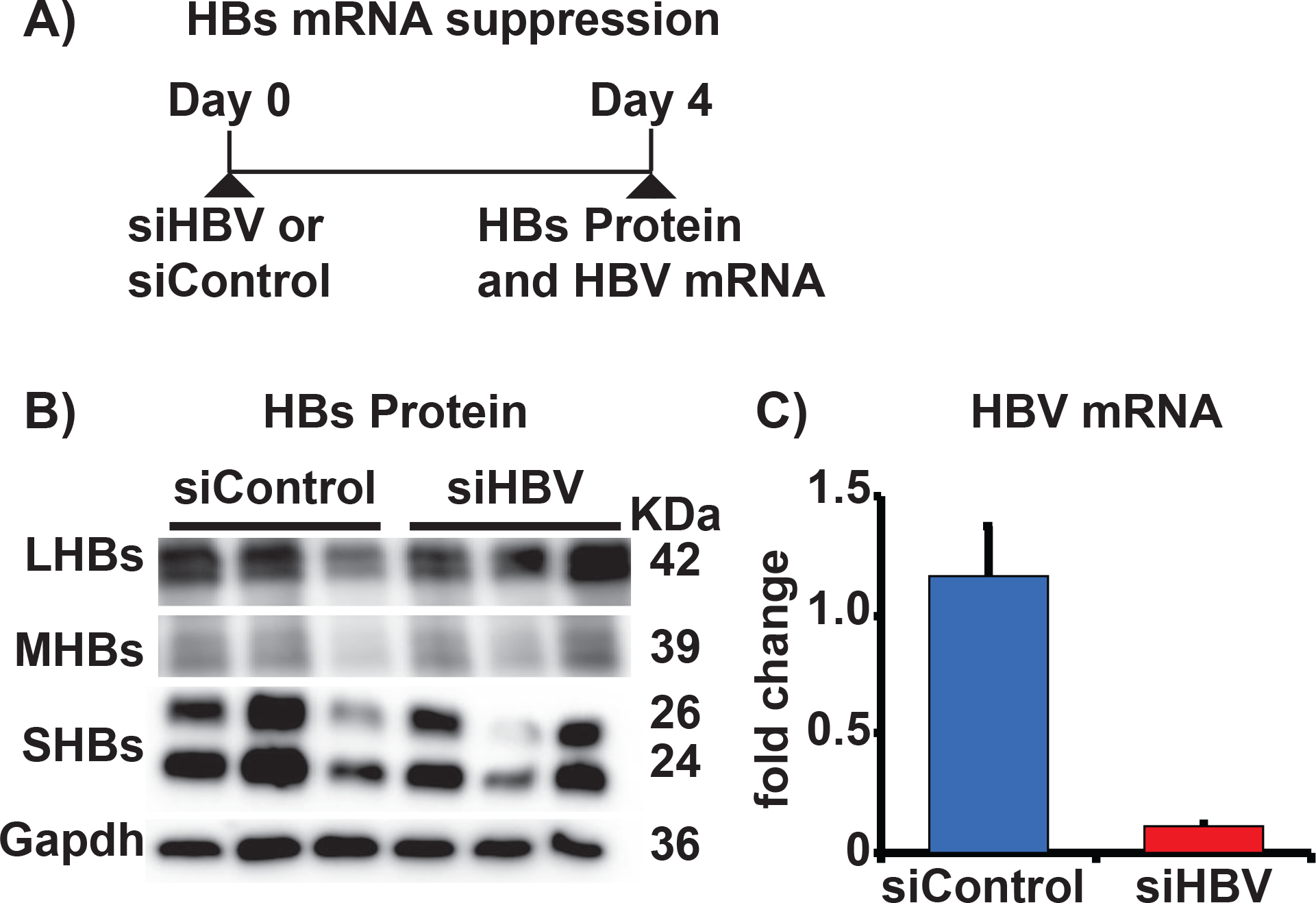
Reduction of intrahepatic HBsAg. (A) Experimental design. In vivo suppression of HBsAg using siRNA. HBs-Tg mice were intravenously injected with mixed siRNA targeting HBV (5mg/kg) then sacrificed at the specified time points. (B) The graph shows >10 fold reduction in HBV mRNA levels by siHBV treatment. (C) HBsAg levels in HBs-Tg mice treated with siHBVcompared with control mice.

**S2 Fig.**
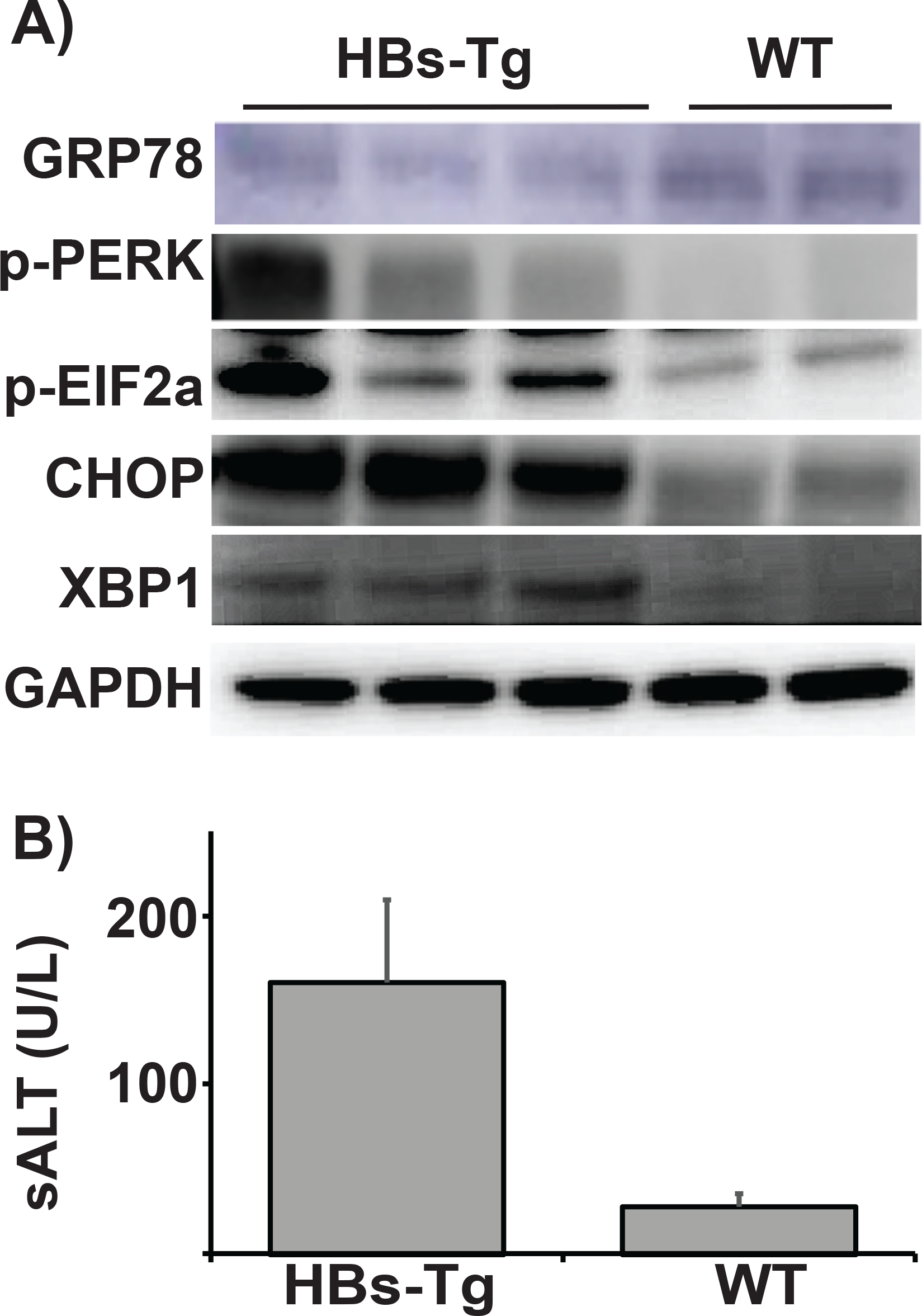
(A) Baseline expression of UPR markers in non-treated 107-5D HBs-Tg mice. The Immunoblots show the expression of UPR molecules like CHOP, XBP1 and GRP78 in non-treated HBs-Tg mice compared with normal WT mice. (B) Serum ALT levels in non-treated HBs-Tg mice compared with normal WT mice.

**S3 Fig.**
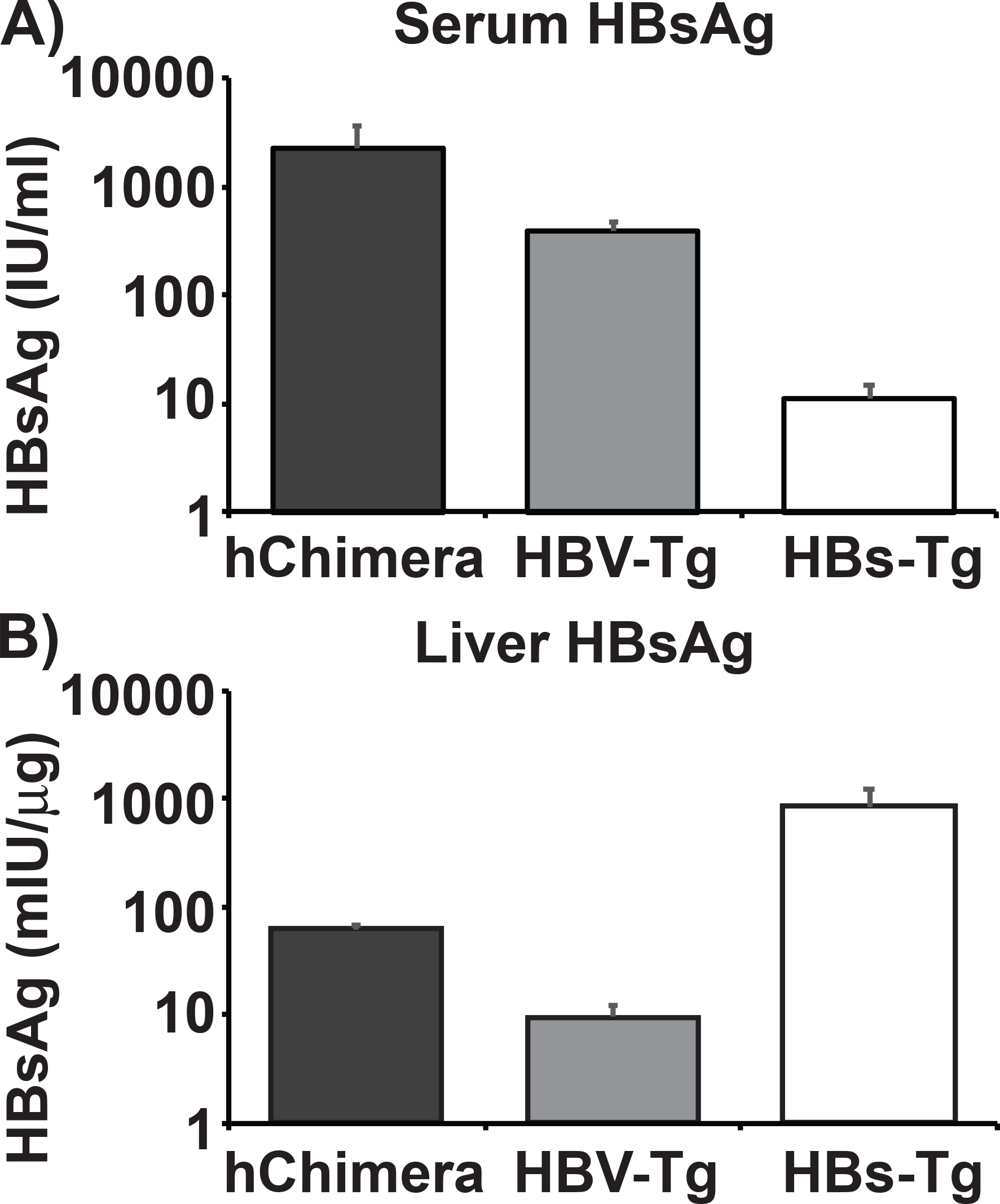
Characterization of the baseline extracellular and intracellular HBsAg levels in non-treated HBV infected chimeric, HBV-Tg and HBs-Tg mice. (A) A graph showing the serum HBsAg levels between chimeric mice, HBV-Tg and HBs-Tg mice. (B) A graph showing the intracellular HBsAg levels per μg of liver protein between chimeric mice, HBV-Tg and HBs-Tg mice.

**S4 Fig.**
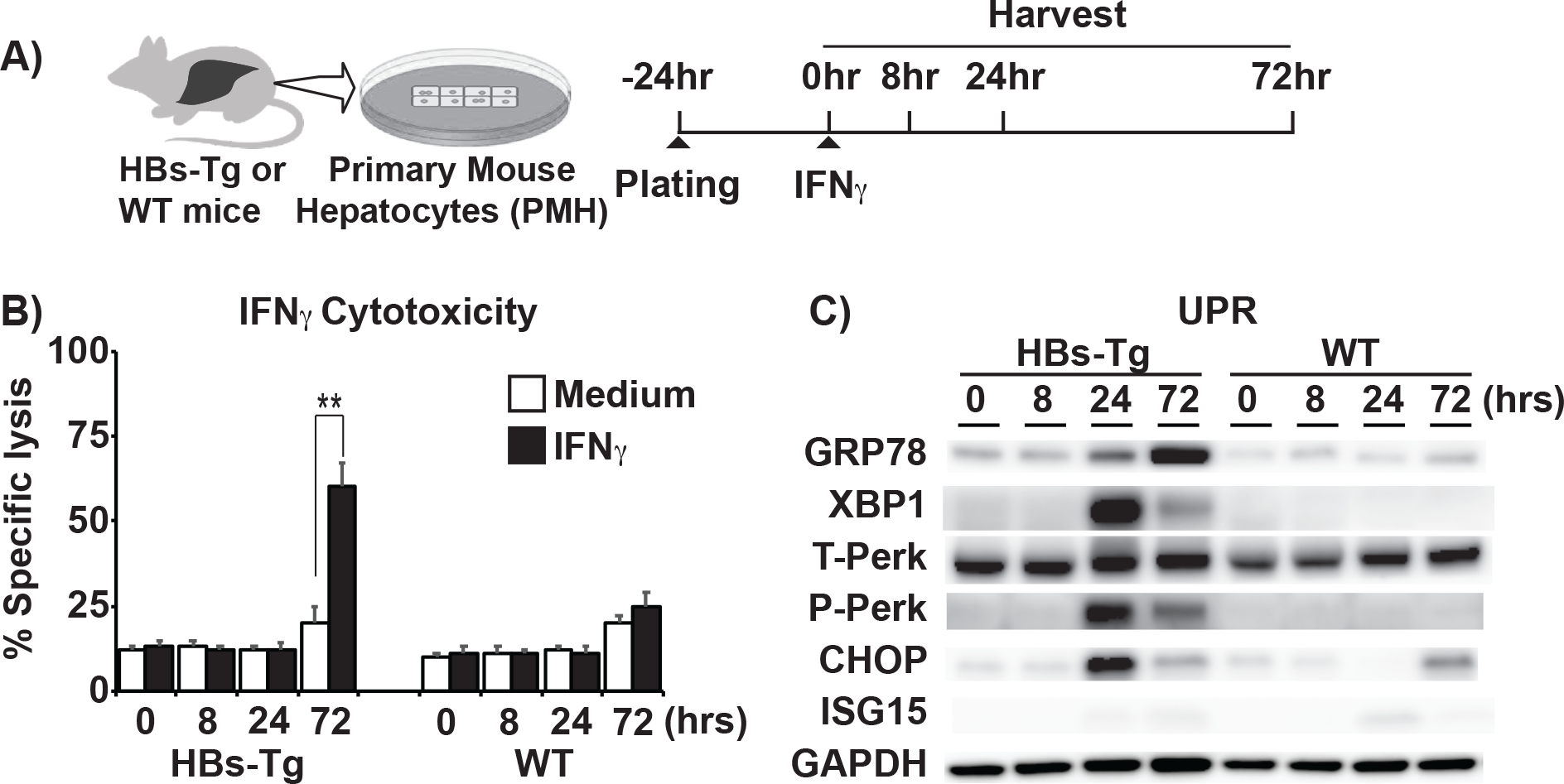
IFNγ is directly cytotoxic to hepatocytes accumulating HBsAg. (A) Timeline showing how primary mouse hepatocytes (PMHs) from both HBs-Tg and WT mice were isolated, plated, and treated with either IFNγ, while monitoring cytotoxicity at the specified time points. (B) The graph shows significant LDH increase in HBs-Tg derived PMHs 72 hours after IFNγ addition. (C) The immunoblots show the effect of IFN-1 signaling on UPR-related proteins at specified time points after IFNγ treatment.

## References

1. Samuel CE. Antiviral Actions of Interferons. Clin Microbiol Rev. 2001;14: 778–809. doi:10.1128/CMR.14.4.778-809.2001

2. McNab F, Mayer-Barber K, Sher A, Wack A, O’Garra A. Type I interferons in infectious disease. Nat Rev Immunol. 2015;15: 87–103. doi:10.1038/nri3787

3. Stetson DB, Medzhitov R. Type I Interferons in Host Defense. Immunity. 2006;25: 373–381. doi:10.1016/j.immuni.2006.08.007

4. Ron D, Walter P. Signal integration in the endoplasmic reticulum unfolded protein response. Nat Rev Mol Cell Biol. 2007;8: 519–529. doi:10.1038/nrm2199

5. Hetz C. The unfolded protein response: controlling cell fate decisions under ER stress and beyond. Nat Rev Mol Cell Biol. 2012;13: 89–102. doi:10.1038/nrm3270

6. Gerakis Y, Hetz C. Emerging roles of ER stress in the etiology and pathogenesis of Alzheimer‘s disease. 2018;285: 995–1011. doi:10.1111/febs.14332

7. Maiers JL, Malhi H. Endoplasmic Reticulum Stress in Metabolic Liver Diseases and Hepatic Fibrosis. Semin Liver Dis. 2019;39: 235–248. doi:10.1055/s-0039-1681032

8. Lindholm D, Korhonen L, Eriksson O, Kõks S. Recent Insights into the Role of Unfolded Protein Response in ER Stress in Health and Disease. 2017;5: 1–16. doi:10.3389/fcell.2017.00048

9. Wang M, Kaufman RJ. The impact of the endoplasmic reticulum protein-folding environment on cancer development. Nat Rev Cancer. 2014;14: 581–597. doi:10.1038/nrc3800

10. Hetz C, Chevet E, Harding HP. Targeting the unfolded protein response in disease. Nat Rev Drug Discov. 2013;12: 703–19. doi:10.1038/nrd3976

11. Yoshida H. ER stress and diseases. FEBS J. 2007;274: 630–658. doi:10.1111/j.1742-4658.2007.05639.x

12. Walter P, Ron D. The unfolded protein response: From stress pathway to homeostatic regulation. Science (80-). 2011;334: 1081–1086. doi:10.1126/science.1209038

13. Wu J, Kaufman RJ. From acute ER stress to physiological roles of the unfolded protein response. Cell Death Differ. 2006;13: 374–384. doi:10.1038/sj.cdd.4401840

14. Tardif KD, Mori K, Siddiqui A. Hepatitis C Virus Subgenomic Replicons Induce Endoplasmic Reticulum Stress Activating an Intracellular Signaling Pathway. J Virol. 2002;76: 7453–7459. doi:10.1128/jvi.76.15.7453-7459.2002

15. Tardif KD, Mori K, Kaufman RJ, Siddiqui A. Hepatitis C Virus Suppresses the IRE1-XBP1 Pathway of the Unfolded Protein Response. J Biol Chem. 2004;279: 17158– 17164. doi:10.1074/jbc.M312144200

16. Zambrano JL, Ettayebi K, Maaty WS, Faunce NR, Bothner B, Hardy ME. Rotavirus infection activates the UPR but modulates its activity. Virol J. 2011;8: 359. doi:10.1186/1743-422X-8-359

17. Perera N, Miller JL, Zitzmann N. The role of the unfolded protein response in dengue virus pathogenesis. Cellular Microbiology. 2017. p. e12734. doi:10.1111/cmi.12734

18. Blázquez AB, Escribano-Romero E, Merino-Ramos T, Saiz JC, Martín-Acebes MA. Stress responses in flavivirus-infected cells: Activation of unfolded protein response and autophagy. Frontiers in Microbiology. 2014. pp. 1–7. doi:10.3389/fmicb.2014.00266

19. Li S, Ye L, Yu X, Xu B, Li K, Zhu X, et al. Hepatitis C virus NS4B induces unfolded protein response and endoplasmic reticulum overload response-dependent NF-κB activation. Virology. 2009;391: 257–264. doi:10.1016/j.virol.2009.06.039

20. Davies SE, Portmann BC, O′grady JG, Aldis PM, Chaggar K, Alexander GJM, et al. Hepatic histological findings after transplantation for chronic hepatitis B virus infection, including a unique pattern of fibrosing cholestatic hepatitis. Hepatology. 1991;13: 150–157. doi:10.1002/hep.1840130122

21. Lau JYN, Bain VG, Davies SE, O’Grady JG, Alberti A, Alexander GJM, et al. Highlevel expression of hepatitis B viral antigens in fibrosing cholestatic hepatitis. Gastroenterology. 1992;102: 956–962. doi:https://doi.org/10.1016/0016-5085(92)90182-X

22. Lok ASF, McMahon BJ. Chronic hepatitis B. Hepatology. 2007. doi:10.1002/hep.21513

23. Dunn C, Brunetto M, Reynolds G, Christophides T, Kennedy PT, Lampertico P, et al. Cytokines induced during chronic hepatitis B virus infection promote a pathway for NK cell–mediated liver damage. J Exp Med. 2007;204: 667–680. doi:10.1084/jem.20061287

24. Tan AT, Koh S, Goh W, Zhe HY, Gehring AJ, Lim SG, et al. A longitudinal analysis of innate and adaptive immune profile during hepatic flares in chronic hepatitis B. J Hepatol. 2010;52: 330–339. doi:10.1016/j.jhep.2009.12.015

25. Deng G, Zhou G, Zhang R, Zhai Y, Zhao W, Yan Z, et al. Regulatory Polymorphisms in the Promoter of CXCL10 Gene and Disease Progression in Male Hepatitis B Virus Carriers. Gastroenterology. 2008;134: 716–726. doi:10.1053/j.gastro.2007.12.044

26. Wong GLH, Chan HLY, Chan HY, Tse CH, Chim AML, Lo AOS, et al. Serum interferon-inducible protein 10 levels predict hepatitis B s antigen seroclearance in patients with chronic hepatitis B. Aliment Pharmacol Ther. 2016;43: 145–153. doi:10.1111/apt.13447

27. Chisari F V., Filippi P, Buras JON, McLachlan A, Popper H, Pinkert CA, et al. Structural and pathological effects of synthesis of hepatitis B virus large envelope polypeptide in transgenic mice. Proc Natl Acad Sci. 1987;84: 6909–6913. doi:10.1073/pnas.84.19.6909

28. Guidotti LG, Matzke B, Schaller H, Chisari F V. High-level hepatitis B virus replication in transgenic mice. J Virol. 1995;69: 6158–69. Available: http://www.ncbi.nlm.nih.gov/pubmed/7666518%0Ahttp://www.pubmedcentral.nih.gov/articlerender.fcgi?artid=PMC189513

29. Yamamoto N, Sato Y, Munakata T, Kakuni M, Tateno C, Sanada T, et al. Novel pH-sensitive multifunctional envelope-type nanodevice for siRNA-based treatments for chronic HBV infection. J Hepatol. 2016;64: 547–555. doi:10.1016/j.jhep.2015.10.014

30. Wooddell CI, Rozema DB, Hossbach M, John M, Hamilton HL, Chu Q, et al. Hepatocyte-targeted RNAi therapeutics for the treatment of chronic hepatitis B virus infection. Mol Ther. 2013;21: 973–985. doi:10.1038/mt.2013.31

31. Ando K, Moriyama T, Guidotti LG, Wirth S, Schreiber RD, Schlicht HJ, et al. Mechanisms of class I restricted immunopathology. A transgenic mouse model of fulminant hepatitis. J Exp Med. 1993;178: 1541–54. doi:10.1084/jem.178.5.1541

32. Conesa A, Talón M, Nueda MJ, Ferrer A. maSigPro: a method to identify significantly differential expression profiles in time-course microarray experiments. Bioinformatics. 2006;22: 1096–1102. doi:10.1093/bioinformatics/btl056

33. Sugiyama M, Tanaka Y, Kato T, Orito E, Ito K, Acharya SK, et al. Influence of hepatitis B virus genotypes on the intra- and extracellular expression of viral DNA and antigens. Hepatology. 2006;44: 915–924. doi:10.1002/hep.21345

34. Gilles PN, Guerrette DL, Ulevitch RJ, Schreiber RD, Chisari F V. HBsAg retention sensitizes the hepatocyte to injury by physiological concentrations of interferon-γ. Hepatology. 1992;16: 655–663. doi:10.1002/hep.1840160308

35. Ito K, Qin Y, Guarnieri M, Garcia T, Kwei K, Mizokami M, et al. Impairment of Hepatitis B Virus Virion Secretion by Single-Amino-Acid Substitutions in the Small Envelope Protein and Rescue by a Novel Glycosylation Site. J Virol. 2010;84: 12850– 12861. doi:10.1128/JVI.01499-10

36. Ou JH, Rutter WJ. Regulation of secretion of the hepatitis B virus major surface antigen by the preS-1 protein. J Virol. 1987;61: 782–786.

37. Ito H, Ando K, Ishikawa T, Saito K, Takemura M, Imawari M, et al. Role of TNF-Produced by Nonantigen-Specific Cells in a Fulminant Hepatitis Mouse Model. J Immunol. 2009;182: 391–397. doi:10.4049/jimmunol.182.1.391

38. Chen Y, Sun R, Jiang W, Wei H, Tian Z. Liver-specific HBsAg transgenic mice are over-sensitive to Poly(I:C)-induced liver injury in NK cell- and IFN-gamma-dependent manner. J Hepatol. 2007;47: 183–190. doi:10.1016/j.jhep.2007.02.020

39. Baerwald KD, Corbin JG, Popko B. Major histocompatibility complex heavy chain accumulation in the endoplasmic reticulum of oligodendrocytes results in myelin abnormalities. J Neurosci Res. 2000;59: 160–169. doi:10.1002/(SICI)1097-4547(20000115)59:2<160::AID-JNR2>3.0.CO;2-K

40. Harding HP, Zhang Y, Bertolotti A, Zeng H, Ron D. Perk is essential for translational regulation and cell survival during the unfolded protein response. Mol Cell. 2000;5: 897–904. doi:10.1016/S1097-2765(00)80330-5

41. Yoshida H, Matsui T, Yamamoto A, Okada T, Mori K. XBP1 mRNA is induced by ATF6 and spliced by IRE1 in response to ER stress to produce a highly active transcription factor. Cell. 2001;107: 881–891. doi:10.1016/S0092-8674(01)00611-0

42. Teske BF, Wek SA, Bunpo P, Cundiff JK, McClintick JN, Anthony TG, et al. The eIF2 kinase PERK and the integrated stress response facilitate activation of ATF6 during endoplasmic reticulum stress. Mol Biol Cell. 2011;22: 4390–4405. doi:10.1091/mbc.e11-06-0510

43. Liu X, Henkel AS, LeCuyer BE, Schipma MJ, Anderson KA, Green RM. Hepatocyte X-box binding protein 1 deficiency increases liver injury in mice fed a high-fat/sugar diet. Am J Physiol Gastrointest Liver Physiol. 2015;309: G965–74. doi:10.1152/ajpgi.00132.2015

44. Olivares S, Henkel AS. Hepatic Xbp1 Gene Deletion Promotes Endoplasmic Reticulum Stress-induced Liver Injury and Apoptosis *. 2015;290: 30142–30151. doi:10.1074/jbc.M115.676239

45. Bertolotti A., Zhang Y., Hendershot L. HH and RD (2000). Dynamic interaction of BiP and the ER stress transducers in the unfolded protein response. Nature Cell Biol. 2, 326–332..pdf. 2000;2.

46. Gething M-J. Role and regulation of the ER chaperone BiP. Semin Cell Dev Biol. 1999;10: 465–472. doi:https://doi.org/10.1006/scdb.1999.0318

47. Fu Y, Wey S, Wang M, Ye R, Liao C, Roy-Burman P, et al. Pten null prostate tumorigenesis and AKT activation are blocked by targeted knockout of ER chaperone GRP78/BiP in prostate epithelium. Proc Natl Acad Sci U S A. 2008;105: 19444–9. doi:10.1073/pnas.0807691105

48. Lee AS. Glucose-regulated proteins in cancer: molecular mechanisms and therapeutic potential. Nat Rev Cancer. 2014;14: 263. Available: https://doi.org/10.1038/nrc3701

49. Burikhanov R, Zhao Y, Goswami A, Qiu S, Schwarze SR. The Tumor Suppressor Par-4 Activates an Extrinsic Pathway for Apoptosis. Cell. 2009;138: 377–388. doi:10.1016/j.cell.2009.05.022

50. Zinszner H, Kuroda M, Wang X, Batchvarova N, Lightfoot RT, Remotti H, et al. CHOP is implicated in programmed cell death in response to impaired function of the endoplasmic reticulum. Genes Dev. 1998;12: 982–995. doi:10.1101/gad.12.7.982

51. Oyadomari S, Koizumi A, Takeda K, Gotoh T, Akira S, Araki E, et al. Targeted disruption of the Chop gene delays endoplasmic reticulum stress-mediated diabetes. J Clin Invest. 2002;109: 525–532. doi:10.1172/JCI200214550

52. Nakagawa H, Umemura A, Taniguchi K, Font-Burgada J, Dhar D, Ogata H, et al. ER Stress Cooperates with Hypernutrition to Trigger TNF-Dependent Spontaneous HCC Development. Cancer Cell. 2014;26: 331–343. doi:10.1016/j.ccr.2014.07.001

53. Wieland S, Thimme R, Purcell RH, Chisari F V. Genomic analysis of the host response to hepatitis B virus infection. Proc Natl Acad Sci U S A. 2004;101: 6669–74. doi:10.1073/pnas.0401771101

54. Perrillo R. Benefits and risks of interferon therapy for hepatitis B. Hepatology. 2009;49: S103–S111. doi:10.1002/hep.22956

55. Liaw Y-F, Chen Y-C, Sheen I-S, Chien R-N, Yeh C-T, Chu C-M. Impact of acute hepatitis C virus superinfection in patients with chronic hepatitis B virus infection. Gastroenterology. 2004;126: 1024–1029. doi:10.1053/j.gastro.2004.01.011

56. Negro F. Hepatitis D Virus Coinfection and Superinfection. Cold Spring Harb Perspect Med. 2014;4: a021550–a021550. doi:10.1101/cshperspect.a021550

57. Sugiyama M, Tanaka Y, Kurbanov F, Maruyama I, Shimada T, Takahashi S, et al. Direct Cytopathic Effects of Particular Hepatitis B Virus Genotypes in Severe Combined Immunodeficiency Transgenic With Urokinase-Type Plasminogen Activator Mouse With Human Hepatocytes. Gastroenterology. 2009;136: 652-662.e3. doi:10.1053/j.gastro.2008.10.048

58. Pollicino T, Cacciola I, Saffioti F, Raimondo G. Hepatitis B virus PreS/S gene variants: Pathobiology and clinical implications. J Hepatol. 2014;61: 408–417. doi:10.1016/j.jhep.2014.04.041

59. Fan YF, Lu CC, Chen WC, Yao WJ, Wang HC, Chang TT, et al. Prevalence and significance of hepatitis B virus (HBV) pre-S mutants in serum and liver at different replicative stages of chronic HBV infection. Hepatology. 2001;33: 277–286. doi:10.1053/jhep.2001.21163

60. Meuleman P, Libbrecht L, Wieland S, De R, Habib N, Kramvis A, et al. Immune Suppression Uncovers Endogenous Cytopathic Effects of the Hepatitis B Virus Immune Suppression Uncovers Endogenous Cytopathic Effects of the Hepatitis B Virus. J Virol. 2006;80: 2797–2807. doi:10.1128/JVI.80.6.2797

61. Bénéchet AP, De Simone G, Di Lucia P, Cilenti F, Barbiera G, Le Bert N, et al. Dynamics and genomic landscape of CD8+ T cells undergoing hepatic priming. Nature. 2019. doi:10.1038/s41586-019-1620-6

62. Isogawa M, Chung J, Murata Y, Kakimi K, Chisari F V. CD40 Activation Rescues Antiviral CD8+ T Cells from PD-1-Mediated Exhaustion. PLoS Pathog. 2013;9: 1–16. doi:10.1371/journal.ppat.1003490

63. Kawashima K, Isogawa M, Hamada-Tsutsumi S, Baudi I, Saito S, Nakajima A, et al. Type I Interferon Signaling Prevents Hepatitis B Virus-Specific T Cell Responses by Reducing Antigen Expression. J Virol. 2018;92. doi:10.1128/jvi.01099-18

